# Themis and Grb2 form a constitutive structural hub in T cell receptor signalling

**DOI:** 10.1101/2025.02.16.638521

**Authors:** Danielle M. Clancy, Alba Sanz-Sanjuan, Elisabeth Gilis, Peter Tougaard, Imke Velghe, Yana Van Droogenbroeck, Jan Felix, Yehudi Bloch, Álvaro Furones Cuadrado, Romain Merceron, Stephan Schenck, Peter Vandenabeele, Janine D. Brunner, Tom Taghon, Dirk Elewaut, Savvas N. Savvides

## Abstract

Positive selection of thymocytes is essential for laying the foundations of the mammalian immune system that include the T cell repertoire, self-tolerance, and prevention of autoimmunity. Themis, the archetypal member of a metazoan protein family featuring distinctive CABIT domains, crucially regulates thymocyte positive selection by linking signalling by the T cell receptor (TCR) to the linker of activation of TCR (LAT). Intriguingly, Themis has been proposed to function via a constitutive complex with the multifunctional adaptor Grb2. Although poised to represent a paradigm shift in our understanding of TCR signalling, the structural and mechanistic basis of such an assembly has remained enigmatic. Here, we present the cryo-EM structure of Themis in complex with Grb2, which reveals how the tandem CABIT domains of Themis engulf the C-terminal SH3 domain of Grb2 (Grb2^SH3C^) to enable its latching onto the proline rich sequence of Themis. The remaining two domains of Grb2 adopt at least three conformational poses set to interact with other binding partners such as Sos1. Structural insights from unbound Themis unmask the pronounced flexibility of the CABIT domains of Themis, which becomes ordered upon binding to Grb2 to create a binding hotspot for their constitutive complex. Indeed, Themis variants that abrogate interactions with Grb2 also fail to activate the tyrosine phosphatase SHP-1 after TCR stimulation, analogous to the functional phenotype of Themis-deficient cells. Collectively, our study draws the blueprint of the Themis-Grb2 complex as a dynamic structural hub in T cell development.

## Introduction

T cells serve as a foundational pillar of the mammalian immune system by mediating critical cellular responses that lead to adaptive immunity. The conversion of thymocytes to effector T cells is controlled via well-regulated developmental, activation and differentiation stages driven by signalling cascades triggered by T cell receptor (TCR) recognition of peptide antigens presented by antigen presenting cells^1,2^. Deficiencies or hyperactivation of T cell development can lead to immunodeficiencies and autoimmune diseases. Thus, improved understanding of TCR signalling in physiology and disease is crucial for the development of immunotherapies and the treatment of immune disorders.

The developmental stages of thymocytes are immunologically well understood based on the presentation of the TCR co-receptors CD4 and CD8 in three distinct stages (double-negative as CD4^−^CD8^−^, double-positive as CD4^+^CD8^+^, single-positive as CD4^+^CD8^−^or CD4^−^CD8^+^)^1,3^. However, the molecular and mechanistic events that steer intracellular signalling following TCR activation remain unclear due to the paucity of structural insights of the intracellular protein complexes involved in these processes. In particular, our structural understanding of the connectivity between the TCR signalling complex and intracellular signalling cascades nucleated by the linker of activation of TCR (LAT) to positive selection of thymocytes is lacking.

Themis (thymocyte-expressed molecule involved in selection), a hitherto structurally uncharacterized protein restricted to the T lineage, has emerged as a critical regulator of thymocyte development^4–10^. Themis features two copies of a novel protein domain termed the CABIT (Cysteine-containing-All-Beta-in-Themis) domain followed by a proline rich sequence (PRS) cassette and an extended disordered C-terminal tail^5^. Genetic deletion of Themis severely impairs development from double-positive thymocytes to mature T cells, resulting in a marked reduction in single-positive thymocytes and peripheral effector T cells^4–8^. Given the exclusive incidence of CABIT domains in metazoans, Themis has served as the prototype of a protein superfamily featuring single or double CABIT domains^5^. Interestingly, the discovery of Themis has led to the identification of other mammalian homologues with distinct expression profiles: Themis2 expressed in B cells, myeloid cells and epithelial cells and the murine-specific intestinal Themis3, as well as the more widely expressed single CABIT domain-containing proteins, GAREM1 and GAREM2^11–13^.

A central aspect to the function of Themis concerns its proposed constitutive association with the ubiquitous cytosolic adapter protein Grb2^14–18^ to mediate regulation of the phosphatase SHP-1 as part of a mechanism that tunes the escalation of signalling after TCR activation^9,19,20^. Thus, the Themis-Grb2 complex has emerged as an immunological nexus poised to connect signalling by the TCR and LAT to guide thymocyte selection. Intriguingly, the idea that Grb2 engages in a long-lived constitutive complex with any protein is unique and arguably at odds with its diverse intracellular roles and multiplicity of binding partners in T cells including the well-known signalling proteins VAV1, CD28, CBL, Sos1, and PLC-γ1, and finally its role in Ras signalling downstream of growth factor receptors^9,10,21–23^. How Grb2 utilizes its dynamic three-domain structure consisting of a central SH2 domain flanked by N- and C-terminal SH3 domains^24–26^ to establish a constitutive complex with the distinctive architecture of Themis in a way that would be compatible with its pleiotropic roles has emerged as a major mechanistic conundrum in T cell signalling.

Here we present the cryogenic electron microscopy (cryo-EM) structure of Themis in complex with Grb2 and show how the tandem CABIT domains of Themis exploit their intrinsic flexibility to tightly grapple onto the C-terminal SH3 domain of Grb2 (Grb2^SH3C^) to potentiate binding of the proline-rich region of Themis to Grb2. In this context, the remaining two domains of Grb2, the central Grb2^SH2^ and the N-terminal Grb2^SH3N^, pivot to at least three conformational poses suggesting their availability for other signalling binding partners as dictated by the interactome of Grb2 as a ubiquitous adaptor protein. By combining these structural insights with an interrogation of the structure-function landscape of the Themis-Grb2 complex we provide the blueprint of a new paradigm in T cell development wherein Themis and Grb2 serve as a constitutive signalling hub that bridges TCR signalling and LAT to steer T lymphocyte maturation.

## RESULTS

### Themis and Grb2 assemble as a stable complex

Since the reporting of the putative constitutive association of Themis with the adaptor Grb2^9,19,20^, direct biochemical and biophysical evidence for this complex had been lacking. The proteolytic susceptibility of full length recombinant Themis produced in HEK293S cells **(Extended Data Fig. 1a)** prompted us to pursue a truncated Themis construct spanning residues 1-563, that lacks the predicted disordered C-terminus but retains the PRS required for Grb2 binding^16^. Recombinant Themis^1–563^ and full-length Grb2 (Grb2^FL^) were produced in HEK293S cells (**Fig. 1a**). Themis^1–563^ forms a high affinity interaction with Grb2^FL^, with an apparent equilibrium dissociation constant (*K*_D_) of 236 nM as determined by isothermal titration calorimetry (ITC) (**Fig. 1b**). The apparent binding in titrations with Themis^1–563^ yielded n-values lower than 1, suggesting that a fraction of recombinant Themis^1–563^ is unavailable for binding, consistent with the observation of a portion of recombinant Themis co-purifying with endogenous cellular Grb2 (**Extended Data Fig. 1b)**.

**Figure 1.**
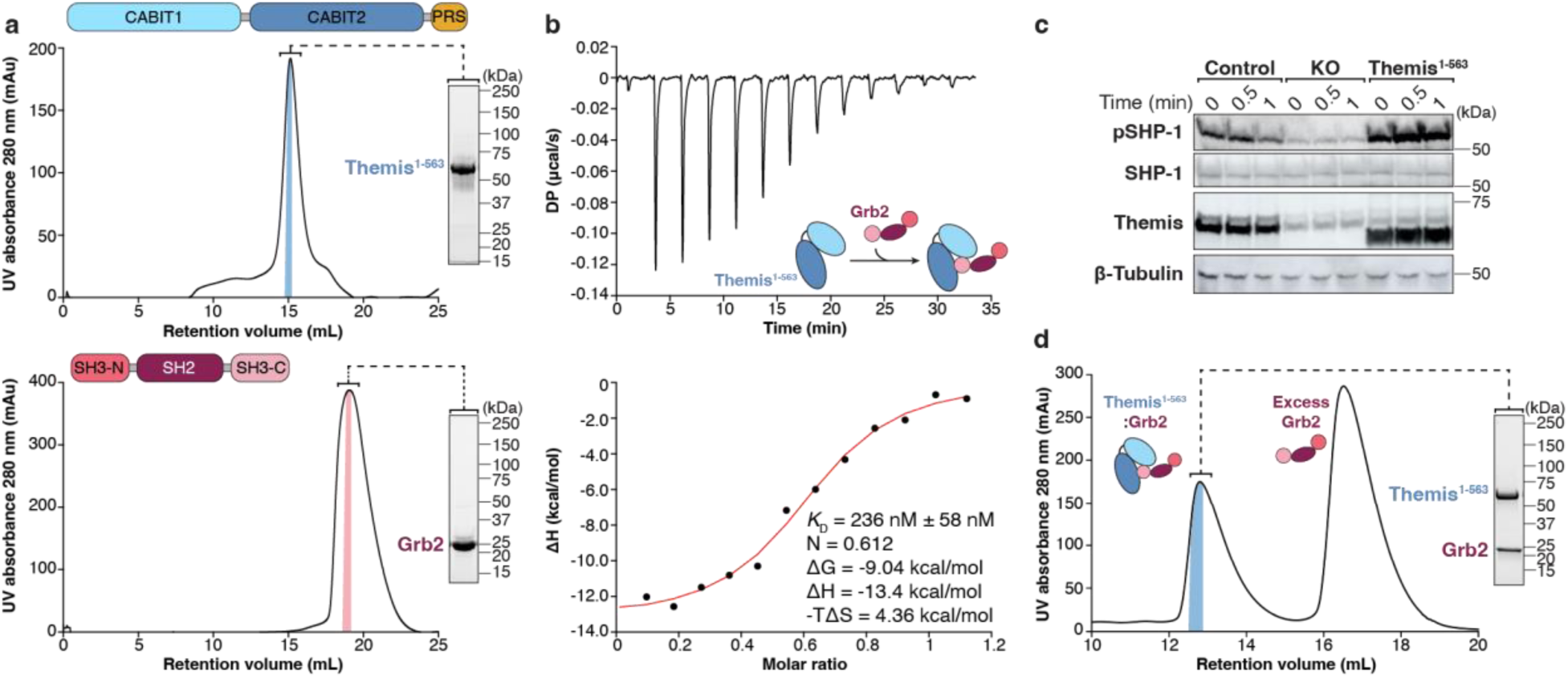
Biochemical and biophysical characterization of the Themis^1-563^-Grb2 complex. **a,** Schematic domain delineations, representative chromatograms and SDS-PAGE gels for the purification of Themis^1-563^ (top) and Grb2^FL^ (bottom). Each protein was purified multiple times and chromatograms and SDS-PAGE analysis are representative for different protein batches. **b,** Representative isothermal titration calorimetry (ITC) thermograms for the titration of Themis^1-563^ (7 μM) with Grb2^FL^ (41.6 μM). DP, differential electrical power. Fitted values are provided with their fitting errors as reported by the MicroCal PEAQ-ITC Analysis software version 1.1.0.1262. Similar results were obtained from four independent experiments. **c,** Western blot analysis of SHP-1 phosphorylation (pY564) after TCR stimulation. Jurkat cells reconstituted with empty lentiviral vector (control), Themis-deficient Jurkat cells (KO) generated by CRISPR/Cas9 gene editing and Jurkat KO cells lentivirally reconstituted with Themis^1-563^ were stimulated with plate-bound anti-CD3 (2 μg/mL) and soluble anti-CD28 (1 μg/mL) for the indicated time. The indicated proteins were analyzed by western blot. Data are representative of three independent experiments. **d,** Representative chromatogram for the co-expression and purification of the Themis^1-563^-Grb2^FL^ complex from HEK293 suspension cells.

Recent studies have suggested that Themis regulates TCR signalling by stabilising SHP-1 in its active state^20^. Under basal conditions, SHP-1 exists in an auto-inhibited state, however TCR activation triggers phosphorylation of the C-terminus of SHP-1, inducing a conformational change that unleashes its phosphatase (PTP) activity^27^. Consistent with this, CRISPR/Cas9-mediated deletion of Themis in Jurkat cells resulted in reduced SHP-1 phosphorylation (pY564) after TCR stimulation (**Fig. 1c**). Reconstitution of Themis knockout (KO) Jurkat cells with a construct encoding the truncated Themis^1–563^ variant restores SHP-1 activation, comparable to endogenous full-length Themis (**Fig. 1c**), indicating that the disordered C-terminal region of Themis is dispensable for its activity downstream of TCR signalling.

### Structure of Themis and architecture of the Themis^1-563^:Grb2 complex

To elucidate the structure of Themis and to generate mechanistic insights into the Themis-Grb2 interaction, we co-expressed Themis^1–563^ and Grb2^FL^ in HEK293S cells. Purified Themis^1–563^– Grb2^FL^ forms a monodisperse 1:1 complex, as demonstrated via size exclusion chromatography coupled to multi-angle laser light scattering (SEC-MALLS, **Fig. 1d** and **Extended Data Fig. 1c**), and was used for structure determination by cryo-EM (**Fig. 2a**). To facilitate structural studies, we utilised a Themis-specific single domain VHH camelid antibody fragment (Nb256). Initial attempts to obtain a high-resolution structure of the Themis^1–563^–Grb2^FL^-Nb256 complex led to 3D model reconstructions up to 4 Å resolution. To overcome this limitation, we employed an engineered nanobody scaffold, termed Pro-Macrobody (PMb), in order to increase the particle size of the Themis^1–563^–Grb2^FL^ complex and to introduce a clear fiducial feature for better particle alignment^28^. To this end, Nb256 was C-terminally fused to a maltose-binding protein (MBP) moiety via a rigid proline linker to create PMb256, which was then complexed with Themis^1–563^–Grb2 (**Fig. 2b and Extended Data Fig. 2a**). The engineered PMb256 retained similar binding affinity and kinetics to Themis^1–563^ compared with Nb256, as measured by biolayer interferometry (BLI) (**Extended Data Fig. 2b,c**).

**Figure 2.**
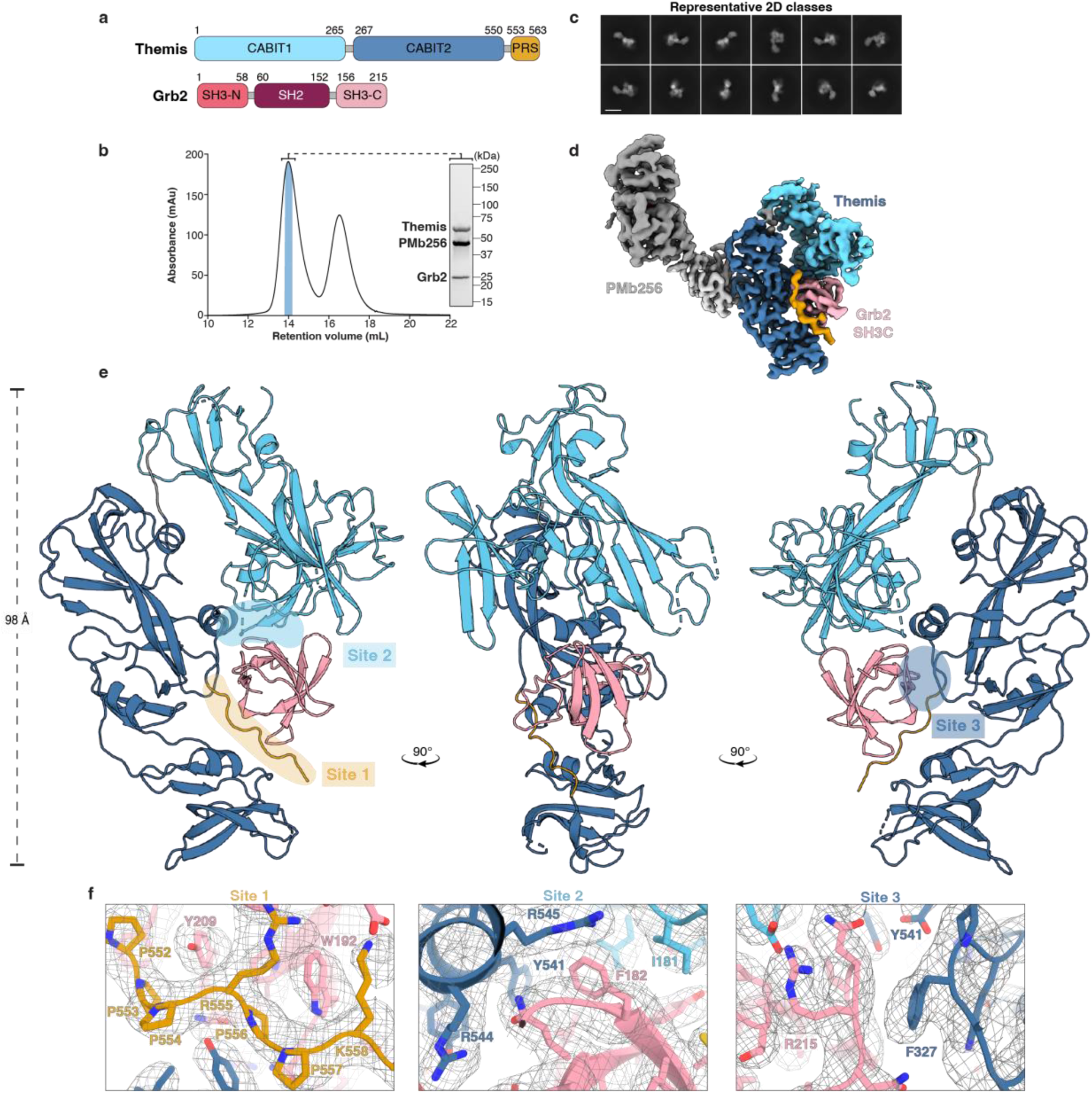
Structure of the Themis^1-563^-Grb2 complex. **a,** Schematic representation of the domain organization of Themis^1-563^ and Grb2. Newly delineated domain boundaries are indicated. **b,** Sample preparation of the Themis^1-563^-Grb2-PMb256 complex for cryo-EM. Representative SEC chromatogram and SDS-PAGE analysis of Themis^1-563^-Grb2 incubated with a 2 molar excess of PMb256. The second SEC peak contains excess PMb256. **c,** Representative cryo-EM 2D class averages of Themis^1-563^-Grb2-PMb256 complex. Scale bars represent 100 Å. **d,** Cryo-EM map of the Themis^1-563^-Grb2-PMb256 complex. The displayed 3D map was sharpened using DeepEMhancer, contoured at 0.12 and coloured per domain with Themis^CABIT1^ domain in cyan, Themis^CABIT2^ in dark blue, Themis^PRS^ in orange, Grb2^SH3C^ in pink and PMb256 in grey. **e,** Real-space refined atomic model of the Themis^1-563^-Grb2^SH3C^ complex. Cartoon representations of front, side and back views are shown, with the PMb256 model removed for clarity. Model is coloured per domain as in **d**. **f,** Zoom-in views of the three interaction sites in the Themis^1-563^-Grb2^SH3C^ interface fitted in a local refinement 3D map displayed as a grey mesh. Grb2^SH3C^, Themis^PRS^, Themis^CABIT1^, Themis^CABIT2^ are shown in pink, orange, cyan and dark blue, respectively.

Cryo-EM analysis of the reconstituted Themis^1–563^–Grb2^FL^-PMb256 complex resulted in a 3D reconstruction to an overall 3.3 Å resolution, and an improved local resolution of 2.8 Å around the Themis^1–563^:Grb2 interaction interface after local refinement (**Fig. 2c,d** ; **Extended Data Fig. 3** and **Table 1**). Model building in the cryo-EM map was achieved using a combination of structural predictions by AlphaFold and ESMFold for Nb256 with an available PMb crystal structure as starting models^28,29^.

**Table 1.**
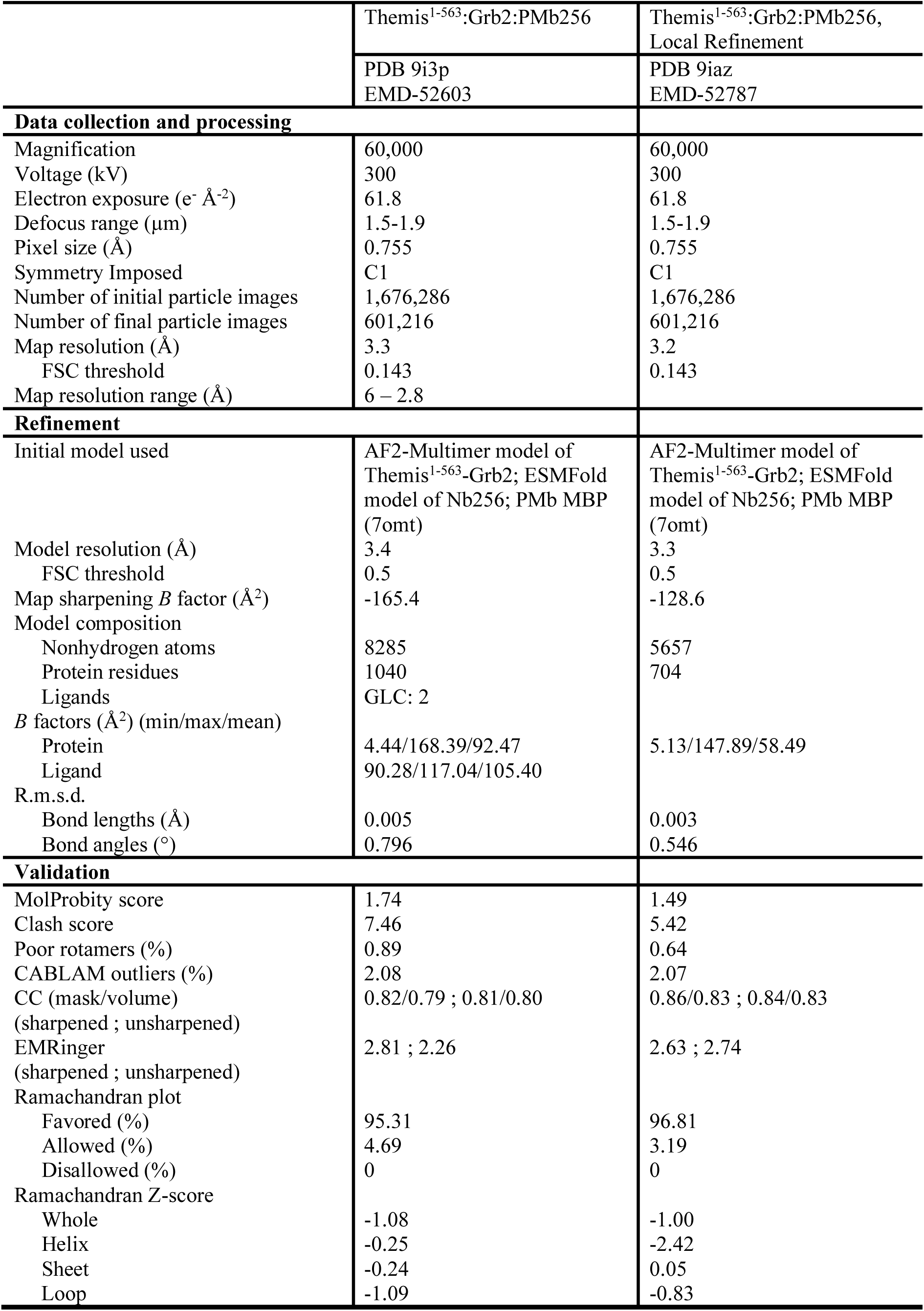
Cryo-EM data collection, refinement, and validation statistics.

|  | Themis<sup>1-563</sup>:Grb2:PMb256 | Themis<sup>1-563</sup>:Grb2:PMb256,<br>Local Refinement |
| --- | --- | --- |
|  | PDB 9i3p<br>EMD-52603 | PDB 9iaz<br>EMD-52787 |
| **Data collection and processing** |  |  |
| Magnification | 60,000 | 60,000 |
| Voltage (kV) | 300 | 300 |
| Electron exposure (e<sup>-</sup> Å<sup>-2</sup>) | 61.8 | 61.8 |
| Defocus range (μm) | 1.5-1.9 | 1.5-1.9 |
| Pixel size (Å) | 0.755 | 0.755 |
| Symmetry Imposed | C1 | C1 |
| Number of initial particle images | 1,676,286 | 1,676,286 |
| Number of final particle images | 601,216 | 601,216 |
| Map resolution (Å) | 3.3 | 3.2 |
| FSC threshold | 0.143 | 0.143 |
| Map resolution range (Å) | 6 – 2.8 |  |
| **Refinement** |  |  |
| Initial model used | AF2-Multimer model of Themis<sup>1-563</sup>-Grb2; ESMFold model of Nb256; PMb MBP (7omt) | AF2-Multimer model of Themis<sup>1-563</sup>-Grb2; ESMFold model of Nb256; PMb MBP (7omt) |
| Model resolution (Å) | 3.4 | 3.3 |
| FSC threshold | 0.5 | 0.5 |
| Map sharpening *B* factor (Å<sup>2</sup>) | -165.4 | -128.6 |
| Model composition |  |  |
| Nonhydrogen atoms | 8285 | 5657 |
| Protein residues | 1040 | 704 |
| Ligands | GLC: 2 |  |
| *B* factors (Å<sup>2</sup>) (min/max/mean) |  |  |
| Protein | 4.44/168.39/92.47 | 5.13/147.89/58.49 |
| Ligand | 90.28/117.04/105.40 |  |
| R.m.s.d. |  |  |
| Bond lengths (Å) | 0.005 | 0.003 |
| Bond angles (°) | 0.796 | 0.546 |
| **Validation** |  |  |
| MolProbity score | 1.74 | 1.49 |
| Clash score | 7.46 | 5.42 |
| Poor rotamers (%) | 0.89 | 0.64 |
| CABLAM outliers (%) | 2.08 | 2.07 |
| CC (mask/volume) | 0.82/0.79 ; 0.81/0.80 | 0.86/0.83 ; 0.84/0.83 |
| (sharpened ; unsharpened) |  |  |
| EMRinger | 2.81 ; 2.26 | 2.63 ; 2.74 |
| (sharpened ; unsharpened) |  |  |
| Ramachandran plot |  |  |
| Favored (%) | 95.31 | 96.81 |
| Allowed (%) | 4.69 | 3.19 |
| Disallowed (%) | 0 | 0 |
| Ramachandran Z-score |  |  |
| Whole | -1.08 | -1.00 |
| Helix | -0.25 | -2.42 |
| Sheet | -0.24 | 0.05 |
| Loop | -1.09 | -0.83 |

The cryo-EM structure presented here reveals that the hitherto uncharacterized CABIT domains of Themis are predominantly composed of beta-sheets, as their name suggests, but also contain multiple short helices (**Fig. 2e**). Previous computational predictions suggested that Themis CABIT domains may adopt a six-stranded beta-barrel fold^5^. Contrary to this, both CABIT domains are composed of two interwoven subdomains, one of which is characterised by two extended beta-sheets, while the other resembles an SH3 domain^5,10^. A recently reported crystal structure for an isolated Themis^CABIT2^ domain is consistent with this fold annotation^30^. The tandem CABIT1 and CABIT2 domains in Themis are structurally distinct and are connected by a short loop. Each CABIT domain contains a highly conserved cysteine residue (Cys152 in CABIT1 and Cys413 in CABIT2)^5,31^ that is buried in the core of the CABIT domain. The CABIT1 and CABIT2 domains were previously predicted to span residues 1-259 and 260-518, respectively^5^ and are currently annotated as such in UniProt^32^. The structural insights revealed here allow us to redefine the boundaries of the Themis CABIT domains. To this end, we now propose that the CABIT1 domain encompasses residues 1-265, while CABIT2 stretches from residues 267-550 (**Fig. 2a**).

Arguably, the hallmark of our structural analysis concerns the way in which Themis and Grb2 engage in the complex. In particular, the structure demonstrates how Themis unexpectedly uses both its CABIT domains to bind to Grb2 via its C-terminal SH3 domain (Grb2^SH3C^) allowing the remaining two domains of Grb2 to be flexible (**Fig. 2e**). We note that PMb256 binds to the CABIT2 domain of Themis^1–563^, distal to the Grb2 interaction interface, and does not interfere with Grb2 binding (**Fig. 2e**).

The core sequence motif of PxRPxK in Themis^PRS^ has previously been implicated as the primary interaction site for Grb2 binding^16^, which we now term site 1. This site 1 interface is mediated by hydrophobic interactions between Themis^PRS^ and conserved aromatic residues in Grb2^SH3C^ (**Fig. 2f** and **Extended Data Fig. 4a**). Unexpectedly, in addition to Themis^PRS^, both CABIT domains of Themis also engage with Grb2^SH3C^. At site 2, a central helix at the end of the Themis^CABIT2^ domain bridges this domain with CABIT1, while projecting the PRS for Grb2^SH3C^ binding. The site 3 interaction interface is driven by interaction of the aromatic residue Themis^F327^ stacking against the main chain of Grb2^R215^ (**Fig. 2f**). Adjacent to site 1, hydrophobic interactions between the CABIT2 SH3-like subdomain and Themis^PRS^ promote PRS stability to facilitate Grb2 binding. Thus, Themis engages Grb2 at three distinct interaction sites, engulfing its SH3C domain to establish a stable complex (**Fig. 2e,f**).

### Both Themis CABIT domains and the PRS synergise to stably bind Grb2

Using structural insights obtained from the Themis^1–563^:Grb2 complex, we interrogated the contribution of the three interaction interfaces involved in Grb2 binding. For mutagenesis studies, we targeted prolines P554 and P557 within the consensus binding motif in Themis^PRS^ at site 1 to verify their role in Grb2 binding. Additionally, we selected residues isoleucine I181, arginine R544 and R545, glutamate E483 and tyrosine Y541 which become buried at the site 2 interface at the junction of the CABIT1 and CABIT2 domains upon Grb2^SH3C^ engagement, as well as phenylalanine F327 at the site 3 interface (**Fig. 2f** and **Extended Data Table 1**). Sequence alignments reveal that the selected residues for mutagenesis are highly conserved across Themis orthologs, suggesting their functional importance in mediating complex formation with Grb2 (**Extended Data Fig. 5**). Interestingly, Grb2 has also been reported to constitutively bind to the related CABIT domain-containing family member, Themis2^14,33^, where the majority of these interfacing residues are also conserved. This suggests that a similar structural mechanism may drive the Themis2-Grb2 interaction (**Extended Data Fig. 6**).

We expressed and purified mutant variants of Themis^1–563^ in which the selected amino acids were mutated to alanine. The biochemical behaviour and stability of all purified mutant Themis variants were comparable to wild-type (WT) Themis (**Extended Data Fig. 7a**). To evaluate the effect of these mutations in binding to Grb2, we employed bio-layer interferometry (BLI). The affinity of the WT Themis^1–563^ interaction with Grb2 as measured by BLI was consistent with the affinity determined by ITC (**Fig. 3a**). Mutations at the site 1 interface, Themis^P554A^ and Themis^P557A^, showed substantially reduced, yet measurable affinity to Grb2 (**Extended Data Fig. 7b**). In contrast, a double mutant Themis^P554A/P557A^ was unable to engage with Grb2, highlighting the critical contribution of the PxRPxK motif in this interaction (**Fig. 3a**). Unexpectedly, mutagenesis at site 2 identified two hotspot residues essential for the Themis:Grb2 interaction. Single point mutants Themis^R544A^ and Themis^R545A^ completely abolished binding to Grb2, exhibiting a greater defect compared to single mutations at the site 1 interface. Similarly, a site 2 double mutant Themis^I181A/R545A^ was also unable to bind Grb2 (**Fig. 3a, Extended Data Fig. 7b**). The remaining site 2 mutations (I181A, E483A, Y541A) displayed reduced binding compared to WT, however the affinity and kinetics of these variants could not be accurately quantified due to poor data fitting (**Extended Data Fig. 7b**). In contrast, binding of Themis^F327A^ was comparable to Themis^WT^, indicating that the site 3 interface is not essential for the Grb2 interaction (**Fig. 3a**). In addition, we also expressed and purified Grb2 variants carrying point mutations at the site 1 and 2 interfaces (**Extended Data Fig. 4b,c**). All Grb2 mutants tested showed either substantially reduced or completely diminished binding to Themis^1-563^ (**Extended Data Fig. 4d**).

**Figure 3.**
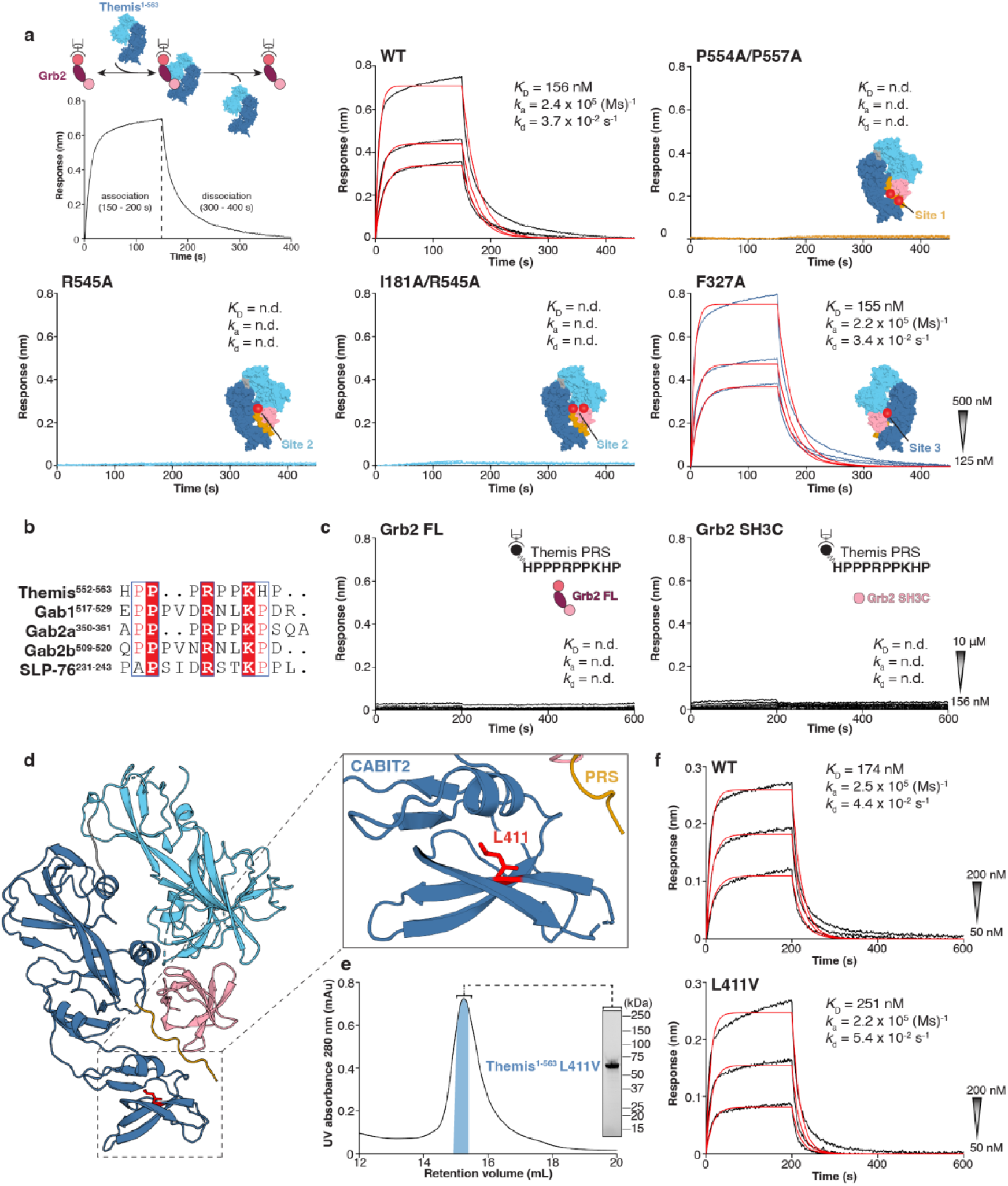
Structural and functional interrogation of Themis^1-563^ mutants. **a,** Experimental BLI setup and representative response curves fitted with a 1:1 binding model (red) to quantify the kinetics (*k*_a_, *k*_d_) and binding affinity (*K*_D_) of wild-type and mutant Themis^1-563^ to Grb2^FL^. The location of the mutations in the Themis^1-563^-Grb2^SH3C^ interface are indicated on a surface representation of the complex. **b,** Sequence alignment of Themis^PRS^ with proline rich motifs from other Grb2^SH3C^ binding partners Gab1, Gab2 and SLP-76. **c,** Representative BLI response curves to quantify the interaction of Grb2^FL^ and Grb2^SH3C^ to an immobilised biotinylated Themis^PRS^ peptide. The peptide sequence is shown inset. **d,** Zoom-in view of the Themis^CABIT2^ domain showing residue L411V coloured in red. **e,** SEC chromatogram and SDS- PAGE analysis for the purification of the Themis^L411V^ mutant variant. **f,** Representative BLI response curves fitted with a 1:1 binding model (red) to compare the kinetics (*k*_a_, *k*_d_) and binding affinity (*K*_D_) of Themis^WT^ and Themis^L411V^ to immobilised Grb2. For all BLI experiments, start and end concentrations of the 2-fold dilution series is shown as an inset.

The identification of binding hotspots in site 2 led us to further investigate the relative importance of the site 1 and site 2 interfaces delineated in the Themis:Grb2 structure using a synthesised biotinylated Themis^PRS^ peptide. Previous studies have reported the specific binding of Grb2^SH3C^ to atypical PxxxRxxKP motifs in the Grb2 binding partners Gab1, Gab2 and SLP-76 (**Fig. 3b**)^34,35^. Notably, Themis^PRS^ deviates from the previously reported Grb2^SH3C^ consensus motif by incorporating a histidine in place of the second conserved proline residue (Fig. 3b). Surprisingly, neither Grb2^FL^ nor the Grb2^SH3C^ domain alone displayed any detectable binding to the Themis^PRS^ peptide, even at high protein concentrations (**Fig. 3c**). In contrast, the isolated Grb2^SH3C^ domain displayed a similar binding affinity to Themis^1-563^ as Grb2^WT^ (**Extended Data Fig. 7c**) Thus, Themis^PRS^ alone is not sufficient for Grb2^SH3C^ binding and instead synergises with the CABIT1 and CABIT2 domains to facilitate stable complex formation.

Interestingly, a recent study identified Themis^L411V^ as a possible pathogenic mutation in rheumatoid arthritis (RA) by whole exome sequencing^36^. Our cryo-EM model now allows structural mapping of this residue and reveals that L411 buried within the SH3-like subdomain of CABIT2 in close proximity to Themis^PRS^ (**Fig. 3d**). Thus, we characterised the Themis^L411V^ variant and found that it displayed comparable behaviour to Themis^WT^ in terms of protein expression, stability and ability to bind Grb2 (**Fig. 3e,f**). To complete our understanding of this reported variant, we investigated its potential pathogenic effect in RA in a transgenic mouse model. T cell development in Themis^L411V^ transgenic mice was comparable to wild-type (WT) littermates, in stark contrast to the significantly reduced numbers of single-positive CD4^+^ and CD8^+^ T cells in the thymus and spleen of Themis^KO^ mice, as previously reported (**Extended Data Fig. 7d**)^4–8^. To evaluate the direct role of Themis^L411V^ in RA, we utilised an experimental collagen-induced arthritis model. Themis^L411V^ mice developed clinical and histopathological signs of arthritis at the same frequency and severity as WT mice, with similar levels of anti-type II collagen IgG1 and IgG2a serum antibodies **(Extended Data Fig. 7e-g).** These results indicate that the L411V mutation does not significantly alter susceptibility to RA and suggests that Themis^L411V^ may require co-segregating pathogenic variants in immune related genes, such as CD36, to contribute to autoimmune disease^36^.

### Themis CABIT domains are highly flexible and become ordered upon Grb2 binding

Whereas our cryo-EM structure of the Themis-Grb2 complex provided unprecedented structural and mechanistic insights, we sought to obtain additional insights from structural studies of Themis in its Grb2-free state by cryo-EM ( **Fig. 4a ; Extended Data Fig. 8**).

**Figure 4.**
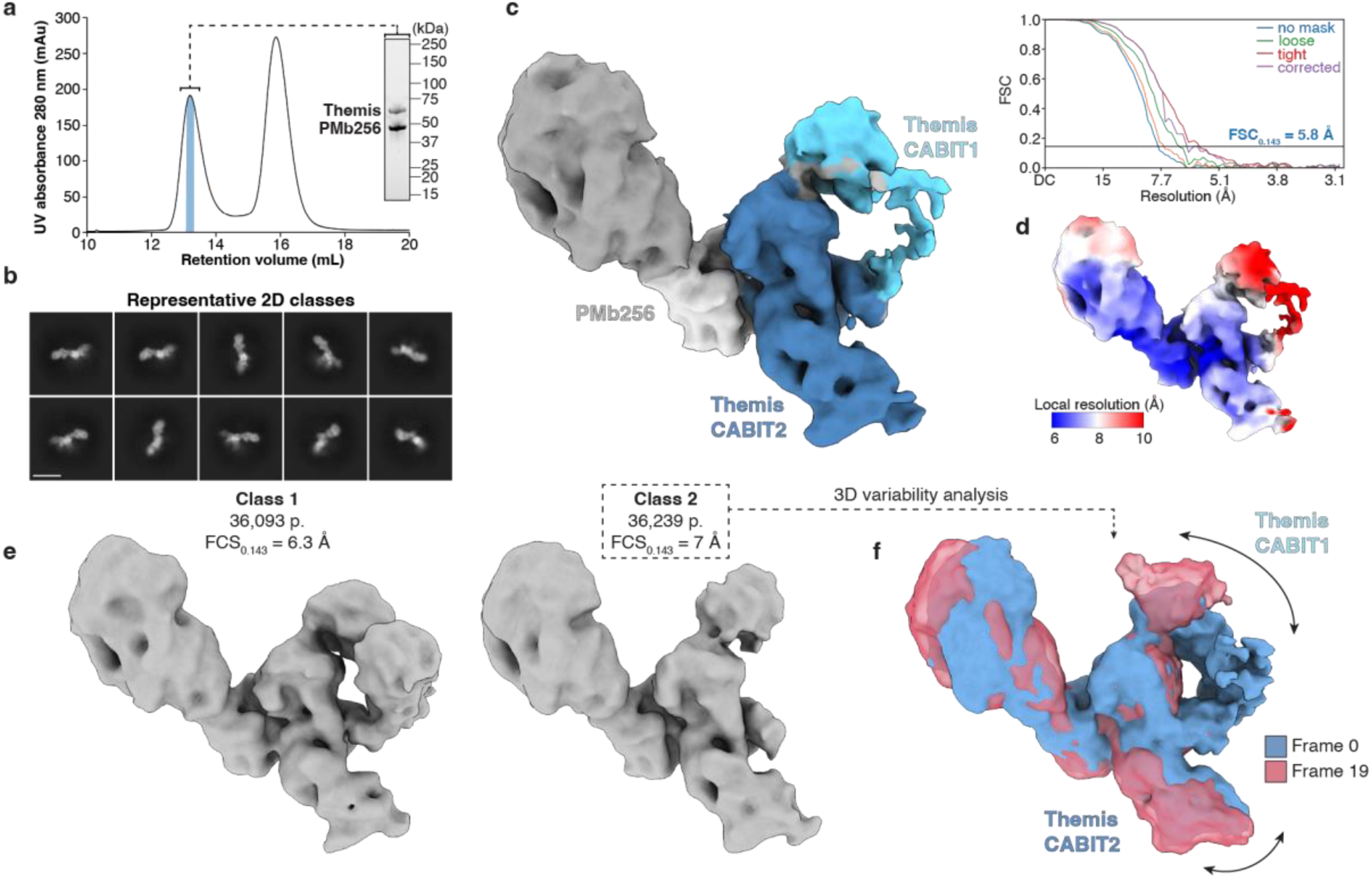
Themis CABIT domains are highly flexible and are stabilized by Grb2 binding. **a,** Sample preparation of the Themis^1-563^-PMb256 complex for cryo-EM. Representative SEC chromatogram and SDS-PAGE analysis of Themis^1-563^ incubated with a 2 molar excess of PMb256. The second SEC peak contains excess PMb256. **b,** Representative cryo-EM 2D class averages of Themis^1-563^-PMb256 complex. Scale bar represents 100 Å. **c,** Cryo-EM map of the Themis^1-563^-PMb256 complex. The displayed 3D map is contoured at 0.12 and coloured with Themis CABIT domains in blue, and PMb256 in grey. Gold-standard Fourier Shell Correlation (FSC) curves are shown, with the estimated resolution at FSC = 0.143 (dark blue line) shown for the corrected FSC curve (purple line). **d,** 3D cryo-EM map shown in **c** colored according to local resolution. **e,** 3D classification without alignment of the particles contributing to the map shown in **c** results in two classes in which the CABIT1 domain is present (class 1) or absent (class 2). Sharpened maps from subsequent non-uniform refinement are shown contoured at 0.12. **f,** 3D variability analysis of particles from class 2 shows movement of the unbound Themis^CABIT1^ domain. 3D volumes from the start (Frame 0, blue) and end (Frame 19, red) of the component analysis are shown.

It was immediately apparent that the 2D classification of particles corresponding to the unbound state of Themis^1-563^ were markedly different from those obtained for the Themis^1-563^- Grb2 complex, with particles appearing more elongated (**Fig. 4b**). The ensuing *Ab-Initio* 3D reconstruction followed by 3D refinement for Themis^1-563^ led to a low resolution map in which the CABIT1 domain is not fully visible (**Fig. 4c**). The lower resolution of this map likely reflects the conformational heterogeneity of Themis in its Grb2-free state, with local resolution estimation indicating 8-10 Å resolution for the CABIT1 domain (**Fig. 4d**). However, this 3D map is reminiscent of the conformational state of the Grb2-bound form of Themis (**Fig. 2e**)

To better investigate the structural plasticity of Themis, we performed 3D classification of the particles contributing to the consensus cryo-EM map and identified two distinct 3D classes, one with evidence for an ordered CABIT1 domain (class 1) and one for a fully disordered CABIT1 domain (class 2) (**Fig. 4e**). Further 3D variability analysis of class 2 revealed extensive movement of the CABIT1 domain, transitioning from a closed state that resembles the conformation of the Grb2-bound protein, to an open state, where the CABIT1 domain extends away from the Grb2 binding site (Fig. 4f). This flexibility is mediated by the short interdomain loop connecting CABIT1 and CABIT2 (residues 265-268) which likely acts as a hinge region allowing movement between these domains. Similarly, the CABIT2 domain also exhibits some conformational flexibility, however it is less pronounced than CABIT1 (**Fig. 4f**), which may be attributable to the binding of PMb256.

The cryo-EM structure of unbound Themis^1-563^ highlights the conformational plasticity and inherent flexibility of the CABIT domains. Grb2^SH3C^ binding anchors Themis by stabilising the central helix mediating the interaction between the CABIT1 and CABIT2 domains, thereby acting as a functional switch to create a structural hotspot for the engagement of Grb2.

### Grb2^SH3N^ and Grb2^SH2^ are flexible in the Themis-Grb2 complex and available for additional partners

A key observation from our model of the Themis^1-563^-Grb2^FL^ complex is that the SH3C domain of Grb2 was the only domain that adopted an ordered structure, serving as the primary interacting domain with Themis (**Fig. 2e and Extended Data Fig. 7c**). This insight suggested that the remaining SH3N and SH2 domains of Grb2 remain flexible, thereby potentially serving to facilitate concomitant interactions with other binding partners along the signalling itinerary of TCR and LAT in thymocyte development. To better understand the flexibility of Grb2 domains in this context, we implemented additional data processing steps to further analyse our structural model. Using a similar approach to that applied to the dataset of unbound Themis, we re-examined the particle set contributing to the final Themis-Grb2 3D reconstruction, aiming to identify additional classes exhibiting discrete conformational heterogeneity.

3D classification analysis revealed four distinct classes with density corresponding to the N-terminal domains of Grb2 (**Fig. 5a**, **Extended Data Fig. 9**). Subsequent processing yielded 3D reconstructions showing at least three distinct conformations of Grb2^SH3N^ and Grb2^SH2^ domains, albeit at low resolution (**Fig. 5b** and **Fig. S9**). Class 4 likely represents an intermediate conformation between Class 2 and Class 3, as indicated by the lower resolution and insufficient density to accommodate both SH2 and SH3N domains using rigid body fitting in ChimeraX.

**Figure 5.**
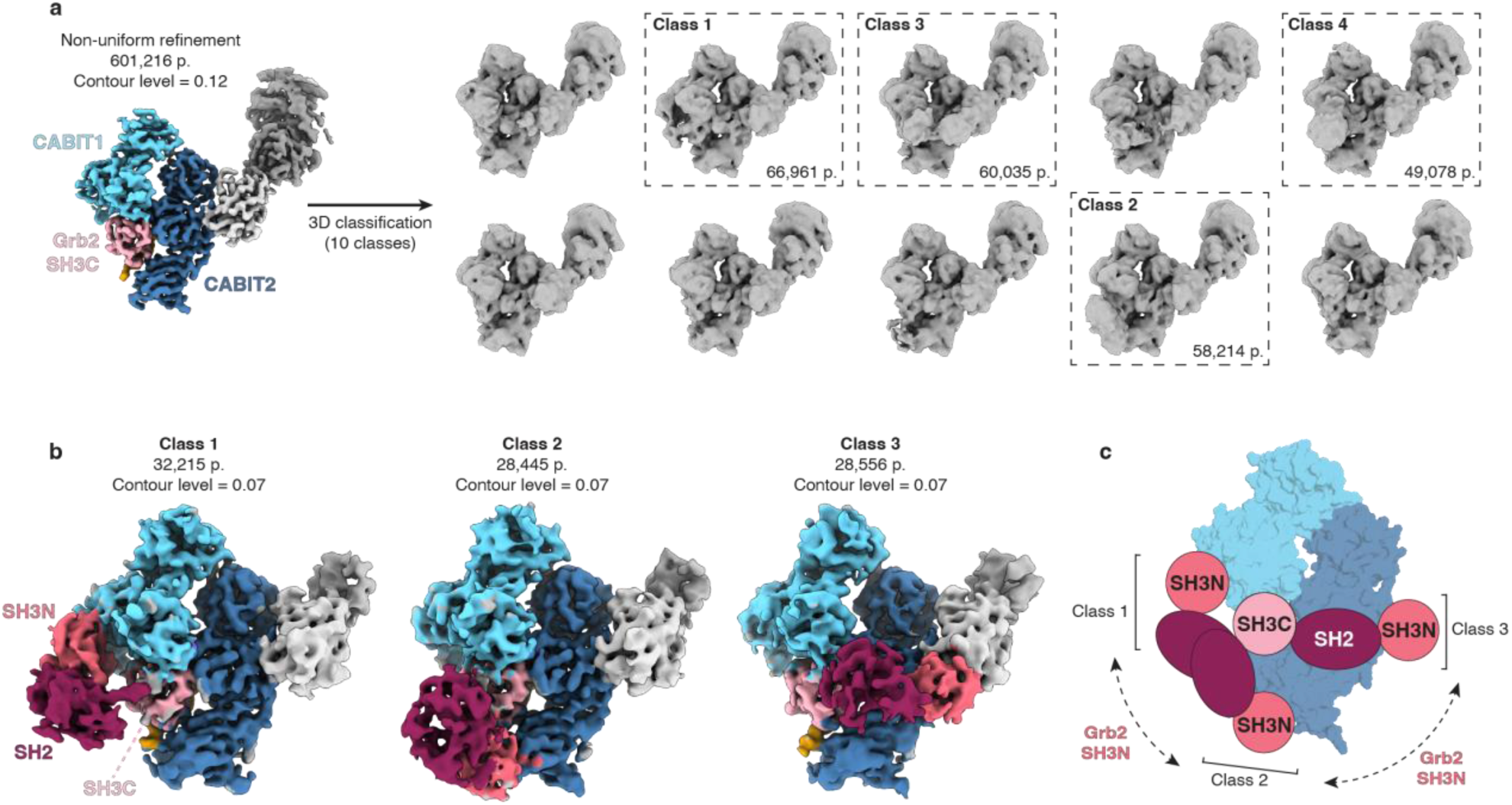
Grb2 SH3N and SH2 domains are highly flexible and sample multiple conformational states. **a,** The particle stack contributing to the consensus map (Fig. 2d) was further processed using 3D classification without alignment in cryoSPARC using 10 classes as input. The resulting classes are shown, with four classes showing volume corresponding to Grb2^SH3N^ and Grb2^SH2^ (dashed boxes). **b,** Maps corresponding to class 1-3 from 3D classification were further processed by particle subtraction to remove the PMb moiety followed by local refinement. The displayed 3D maps are contoured at 0.07 and coloured as before, with the additional Grb2^SH3N^ and Grb2^SH2^ domains in dark pink and wine red, respectively. Particle numbers (p.) contributing to the map for each class are shown. For all maps, the back view of the complex is shown. **c,** Schematic showing the distinct conformational states of Grb2 captured by cryo-EM analysis. Grb2^SH3C^ domain is locked in one position through its interaction with Themis, while Grb2^SH2^ and particularly Grb2^SH3N^ exhibit 180-degree movement at the back of the Themis-Grb2 molecule.

These results demonstrate that while Grb2^SH3C^ remains anchored via its interaction with Themis, Grb2^SH3N^ and Grb2^SH2^ exhibit nearly 180-degree range of motion at the rear of the Themis^1-563^-Grb2 molecule, highlighting the dynamic nature of this complex (**Fig. 5c**). This analysis fortuitously captures the continuum of Grb2 flexibility and suggests the availability of Grb2^SH3N^ and Grb2^SH2^ for interactions with other binding partners after TCR stimulation.

### The Themis:Grb2 complex acts as a signalling hub downstream of TCR activation

The structural model of the Themis^1-563^-Grb2 complex presented here now provides the critical missing link to further explore its role in T cell signalling. Previous studies have established that the Themis-Grb2 interaction is indispensable for recruiting Themis to the transmembrane adapter LAT after TCR activation^16^. We propose that upon recruitment to LAT, the Themis-Grb2 complex acts as a dynamic signalling platform, orchestrating the recruitment of other key binding partners to regulate downstream signal transduction.

As previously shown, Themis is required for SHP-1 phosphorylation in response to TCR engagement^15^ (**Fig. 1c**). To investigate the role of the Themis-Grb2 complex in this pathway, we reconstituted Themis-deficient Jurkat cells with either Themis^WT^ or Themis constructs carrying mutations at the three interaction sites with Grb2^SH3C^, which abrogate complex formation as reported in our interaction studies (**Fig. 3**). Overexpression of Themis^WT^ restores SHP-1 phosphorylation after TCR stimulation, similar to endogenous Themis and previous results with reconstituted Themis^1-563^ (**Fig. 6a** and **Fig. 1c**). Notably, the double mutants at Themis site 1 and site 2 (P554A/P557A and I181A/R545A respectively), which disrupt Grb2 binding, fail to activate SHP-1 after TCR stimulation, exhibiting a similar phenotype to Themis^KO^ cells. Strikingly, a similar result was seen with the single site 2 mutant Themis^R545A^, highlighting this residue as a *bona fide* structural and functional hotspot. The site 3 mutant Themis^F327A^ promotes SHP-1 activation similar to Themis^WT^, confirming the redundancy of site 3 for the Grb2^SH3C^ interaction (**Fig. 6a**). Additionally, we observed no differences in TCR-induced ERK activation between Themis^KO^ and empty vector-transduced control cells, or upon reconstitution of any Themis mutants (**Fig. 6a**). This underscores the specificity of the Themis-Grb2 complex in regulating SHP-1 activation after TCR stimulation.

**Figure 6.**
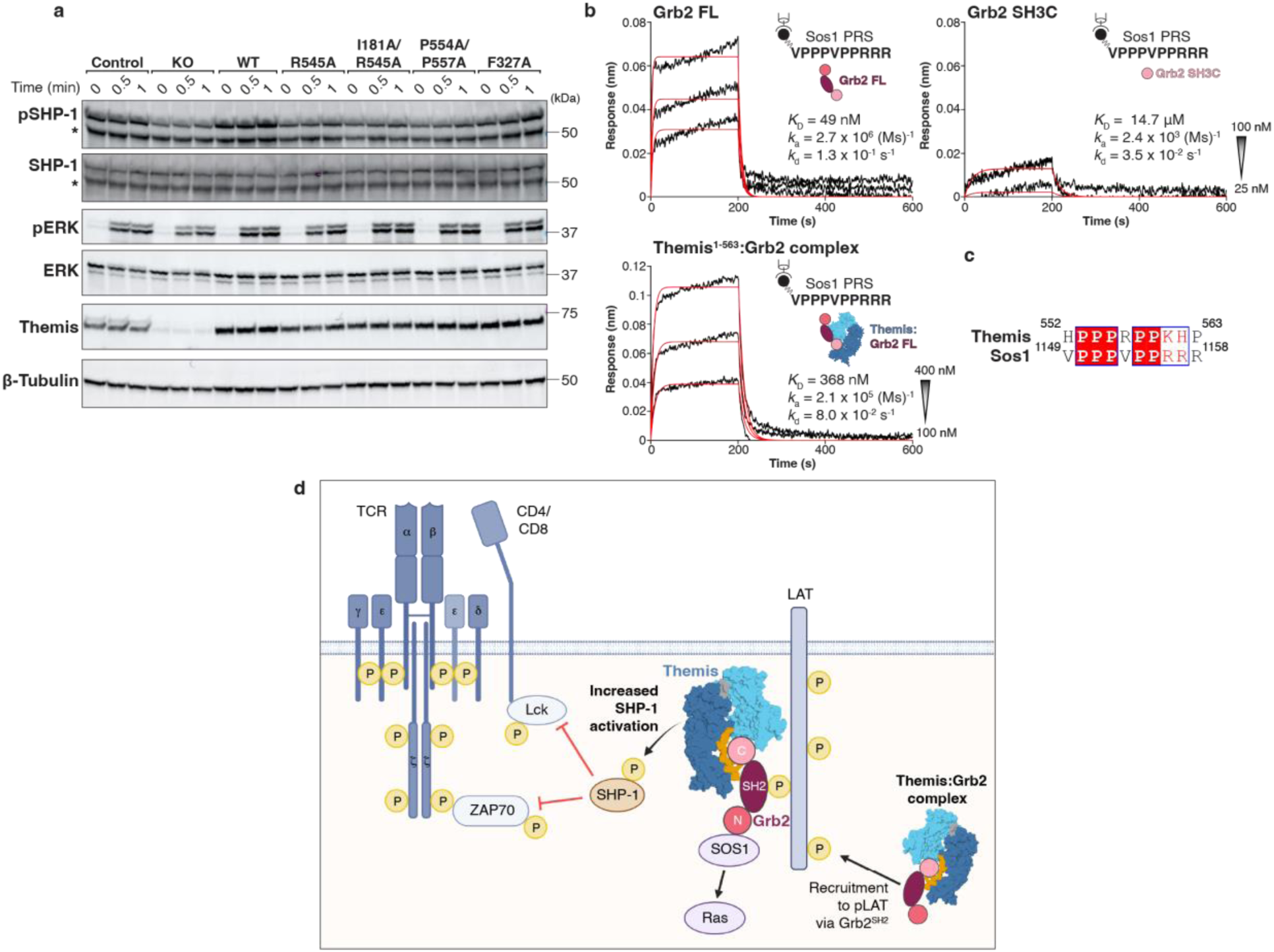
Themis-Grb2 complex acts as a signalling hub downstream of TCR activation. **a,** Western blot analysis of SHP-1 phosphorylation (pY564). Themis-deficient Jurkat cells (KO) were transduced with a lentiviral vector expressing full-length wild-type Themis (WT) or full-length Themis mutants, as indicated. Jurkat cells reconstituted with empty lentiviral vector were used as a control. Cells were stimulated with plate-bound anti-CD3 (2 μg/mL) and soluble anti-CD28 (1 μg/mL). The indicated proteins were analysed by western blot. Asterisks (*) indicate non-specific bands. **b,** Representative response curves fitted with a 1:1 binding model (red) to quantify the kinetics (*k*_a_, *k*_d_) and binding affinity (*K*_D_) of Grb2^FL^, Grb2^SH3C^ and Themis^1-563^-Grb2^FL^ complex to Sos1 PRS peptide. Schematics show the BLI setup for each experiment. Start and end concentrations of the 2-fold dilution series is shown as an inset. **c,** Sequence alignment of the Themis^PRS^ and Sos1^PRS^ peptide. Residue numbers for each peptide are indicated. **d,** Schematic depicting the role of the Themis-Grb2 complex as a proximal signalling hub downstream of the TCR. Upon TCR stimulation, the tyrosine kinase Lck is recruited to the membrane via the CD4 and CD8 co-receptors. Lck phosphorylates immunoreceptor tyrosine-based activation motifs (ITAMs) in the TCR CD3 chains, facilitating the recruitment of the protein tyrosine kinase ZAP70. ZAP70 phosphorylates the transmembrane adapter protein LAT, promoting the formation of the LAT signalosome. The Themis-Grb2 complex is recruited to the membrane via Grb2^SH2^ binding to phosphorylated LAT. Once there, Themis is positioned to interact with the protein tyrosine phosphatase SHP- 1 and regulate its activity to fine-tune TCR signalling. Simultaneously, Grb2^SH3N^ domain is available for binding to other target proteins, contributing to the activation and regulation of other downstream signalling pathways such as the Sos1-Ras-MAPK pathway. Tyrosine phosphorylation (P) is depicted in yellow.

In addition to its interaction with Themis, the role of Grb2 in TCR-induced Ras activation has been well established. In this context, Grb2 plays a similar role in recruiting Son-of-sevenless-homologue 1 (Sos1) to the membrane to promote Ras activation following TCR activation^2,37^. As previously discussed, the SH3 domains of Grb2 have distinct binding preferences to proline rich sequences, with Grb2^SH3N^ preferentially interacting with consensus PxxPxR motifs, as found in Sos1, while Grb2^SH3C^ favors PxRPxK motifs, as found in Themis^34^ (**Fig. 6c** and **Fig. 3b**). Our structural analysis by cryo-EM has uncovered the flexibility and binding availability of the Grb2^SH3N^ domain when Grb2 is bound to Themis^1-563^, leading us to speculate if the Themis-Grb2 complex can engage with additional signalling partners such as Sos1.

To further explore this possibility, we tested a biotinylated Sos1^PRS^ peptide (VPPPVPPRRR) in binding studies by BLI. Sos1^PRS^ binds Grb2^FL^ with high affinity (*K*_D_ = 49 nM), but only weakly binds to Grb2^SH3C^ (*K*_D_ = 14.7 μM), confirming the preferential binding of Sos1^PRS^ to Grb2^SH3N^. Intriguingly, a pre-formed Themis-Grb2 complex also displayed effective binding to Sos1^PRS^, albeit with lower affinity than Grb2^FL^ alone (**Fig. 6b**). These data suggest the possibility of the formation of a Themis-Grb2-Sos1 ternary complex.

The structural and mechanistic insights described here position the Themis-Grb2 complex as a proximal signalling nexus following TCR activation. Upon receptor stimulation, the constitutive Themis-Grb2 complex is recruited to the immunological synapse via Grb2^SH2^- mediated binding to phosphorylated LAT. Once at the membrane, Themis is poised to interact with SHP-1, modulating its phosphatase activity to fine-tune TCR signalling. Concomitantly, Grb2^SH3N^ remains available to engage additional signalling partners, such as Sos1, coordinating the regulation of other downstream signalling pathways (**Fig. 6d**).

## DISCUSSION

Nearly two decades since the identification of Themis as the archetypal member of a new metazoan protein family in thymocyte development, we reveal here the complete architecture of its unprecedented CABIT domains and provide the structural basis of the constitutive Themis-Grb2 complex in TCR signalling. While the essential role of Themis in T cell development is well established, the structural mechanisms underlying this function have remained elusive.

Themis contains two CABIT domains and a C-terminal PRS, all of which are critical for its activity^5,14,31^. The cryo-EM structure presented here shows that Themis CABIT domains are structurally distinct and adopt a novel fold comprising two interwoven subdomains (**Fig. 2**). The structure of unbound Themis highlights the flexibility and dynamic nature of the CABIT domains, particularly CABIT1 (**Fig. 4**). Engagement of Grb2^SH3C^ reduces this flexibility by stabilising the central helix bridging Themis^CABIT1^ and Themis^CABIT2^, thereby projecting Themis^PRS^ for Grb2 binding.

Themis leverages its unique topology to anchor Grb2^SH3C^ and form a stable, constitutive complex. We demonstrate that Themis^PRS^ and both CABIT domains cooperatively engage Grb2^SH3C^ in a high affinity interaction. Strikingly, Themis^PRS^ alone is not sufficient to induce Grb2 binding (**Fig. 3c**), in stark contrast to PRS peptides from other Grb2 partners including Gab1, Gab2, Sos1 and SLP-70^34,35^. Rather, our structural model reveals three interaction interfaces, wherein Themis^PRS^ synergises with the CABIT domains to engulf Grb2^SH3C^, with site 2 containing bona fide interaction hotspot residues critical for binding (**Fig. 3** and **Extended Data Fig. 7**). These structural insights support previous biochemical results showing that Grb2 fails to interact with truncated Themis mutants lacking either CABIT domain^19,31^ and rationalises the long-lived, constitutive nature of this complex.

The structural and functional insights obtained from this study establish the Themis-Grb2 complex as a key proximal assembly platform following TCR engagement, bridging the LAT signalosome to diverse downstream signalling pathways. Upon receptor stimulation, the Themis-Grb2 complex is recruited to phosphorylated LAT at the membrane via Grb2^SH2^, where it is positioned to interact with and regulate SHP-1^20^. Notably, Themis variants incapable of binding Grb2 fail to activate SHP-1 (**Fig. 6a**), underscoring the central role of the Themis-Grb2 complex in TCR-mediated signal transduction. In parallel, Themis itself is rapidly phosphorylated by Lck and ZAP70 following TCR activation^16,20,38^. A recent study identified the highly conserved residue Tyr34 in Themis^CABIT1^ as the primary site of Lck-mediated tyrosine phosphorylation. While the Themis-Grb2 interaction occurs independently of phosphorylation, Tyr34 phosphorylation (pY34) is essential for Themis association with the SHP-1 PTP domain, stabilising SHP-1 in an active state and increasing its phosphatase activity^20^. Moreover, previous studies have proposed the formation of a Themis-Grb2-SHP-1 trimolecular complex, with evidence indicating enhanced interaction between Themis and SHP-1 in the presence of Grb2^19,39^. Indeed, a structural model of the Themis-Grb2-SHP-1 ternary complex predicted by AlphaFold3^29^ supports this hypothesis, with Themis pY34 predicted to interact with the catalytic pocket of the SHP-1 PTP domain, distal to the Grb2^SH3C^ interaction interface (**Extended Data Fig. 10**). While a direct interaction between SHP-1 and Grb2 is not predicted in this model, the increased rigidity of Themis^CABIT1^ induced by Grb2^SH3C^ binding likely stabilises Y34, facilitating the interaction with SHP-1.

A surprising finding from our structural analysis concerned the lack of defined density for the N-terminal domains of Grb2 in the consensus map of the Themis^1-563^-Grb2 complex. Further particle classification revealed extensive flexibility in the unbound domains of Grb2, with Grb2^SH2^ and Grb2^SH3N^ captured in at least three distinct orientations (**Fig. 5**). Previous studies have highlighted the dynamic nature of Grb2, particularly in the linker regions connecting Grb2^SH2^ and both SH3C domains^25,26^. This flexibility allows Grb2 to adjust the positioning of its SH3N and SH3C domains to bivalently interact with additional binding partners within its diverse interactome^25,26,40^. While Themis-bound Grb2 can still interact with a Sos1^PRS^ peptide via its SH3N domain (**Fig. 6b**), the formation of a Themis-Grb2-Sos1 ternary complex, as well as the potential of the Themis-Grb2 complex to cooperate, or indeed compete, with Sos1-driven Ras activation requires further investigation.

Collectively, the structural and mechanistic insights presented here position the Themis-Grb2 complex as a foundational pillar regulating signal transduction during T cell development and provide a structural blueprint of the molecular interactions driving TCR signalling. These findings deepen our understanding of the Themis-Grb2 complex in thymocyte selection, and its pivotal role in regulating SHP-1 phosphatase activity and may fuel further efforts to target this complex for therapeutic purposes. Additionally, related CABIT domain-containing family members, such as Themis2 and the GAREM proteins, have been reported to bind Grb2 downstream of various cell surface receptors including the B-cell and EGF receptors^14,33,41–43^, suggesting a shared function of CABIT domain proteins in assembling regulatory membrane-proximal signalling platforms across diverse receptor pathways. However, further investigation is required to elucidate the structure of these related CABIT family proteins, and to characterise their interactions with binding partners such as Grb2.

## METHODS

### Plasmid generation for protein expression in mammalian cells

Human cDNA sequences were codon optimised for protein expression in mammalian cells and purchased from Integrated DNA Technologies (IDT) as geneblocks. The sequences were cloned into a modified pHLSec vector^44^, in which the secretion signal was removed, containing a C-terminal hexahistidine (His_6_)-tag preceded by a caspase-3 cleavage site (DEVD) for tag removal. All cloning was carried out using a traditional restriction-ligation approach. Restriction enzymes were purchased from New England Biolabs (NEB). Sequences encoding a truncated Themis variant (Uniprot ID Q8N1K5), comprising amino acids 1-563, and full-length Grb2 (Uniprot ID P62993), comprising amino acids 1-217, were used.

### Protein expression in HEK293S and purification from cell lysates

Production of Themis^1-563^ and Grb2^FL^ was performed in suspension-adapted HEK293S cells (obtained from Prof. Nico Callewaert, VIB-UGent Center for Medical Biology, Ghent, Belgium), either alone to isolate individual proteins or co-expressed to isolate the protein complex. Cells were maintained in a 1:1 ratio of Freestyle 293 Expression Medium (Gibco) and Ex-Cell 293 Serum-Free Medium (Merck). Transient transfection was performed in Freestyle medium at a density of 3 x 10^6^ cells mL^-1^, using linear polyethylenimine (PEI) 25 kDa (Polysciences) as transfection reagent and 4.5 μg DNA and 9 μg PEI per milliliter of transfection volume. For co-expression of the Themis^1-563^-Grb2^FL^ complex, equal ratios of plasmid DNA were co-transfected. Five hours post-transfection, an equal volume of Ex-Cell medium supplemented with penicillin/streptomycin (10^6^ units L^-1^ penicillin, 1 g L^-1^ streptomycin, Gibco) was added, resulting in a final cell density of 1.5 x 10^6^ cells mL^-1^.One day post-transfection, valproic acid was added to a final concentration of 3.5 mM, and glucose to a final concentration of 5 mg mL^-1^.

Two days post-transfection, cells were harvested by centrifugation at 400 g for 10 min. Cell culture supernatant was discarded and the remaining cell pellet was resuspended in lysis buffer (25 mM Tris pH 8, 100 mM NaCl, 2 mM DTT, cOmplete EDTA-free protease inhibitor cocktail (Roche)). Cells were lysed by sonication using a Q500 sonicator (QSonica) for 30 s, 60 % amplitude, 8 cycles. The resulting cell lysate was centrifuged at 15,000 g for 30 mins at 4 °C to remove insoluble material and cellular debris. The clarified medium was filtered through a 0.4 μm filter prior to chromatographic purification steps. The C-terminally His-tagged Themis^1-563^ or Grb2^FL^ were captured via immobilized metal affinity chromatography (IMAC) on a cOmplete His-tag Purification Column (Roche). In the case of co-expression of the complex, Grb2^FL^ was His-tagged and co-expressed with a tagless Themis^1-563^ construct and the complex was purified by capturing His-tagged Grb2 on the aforementioned affinity column. The column was washed with purification buffer (25 mM Tris pH 8, 100 mM NaCl, 2 mM DTT) supplemented with 20 mM imidazole, and proteins were eluted in purification buffer supplemented with 250 mM imidazole. Imidazole in the resulting protein fraction was removed using a HiPrep 26/10 Desalting column (Cytiva). The C-terminal His_6_ tag was removed by overnight digestion with caspase 3 (Addgene, #11821) at 20°C. Undigested protein and the His_6_ tagged enzyme were removed by IMAC. The flowthrough from the second IMAC step, containing the cleaved tagless protein was concentrated and injected onto a Superdex 200 increase 10/300 GL column (Cytiva) and fractions containing the proteins of interest were pooled and stored at −80 °C. The purity of all recombinant proteins was evaluated by SDS-PAGE analysis.

GST-tagged Grb2^SH3C^ was expressed and purified as described previously^35^. Briefly, BL21 (DE3) bacteria were grown in lysogeny broth (LB) and protein expression was induced with 0.05 mM isopropyl β- d-1-thiogalactopyranoside (IPTG) (Goldbio) at 18 °C, overnight. Bacteria were harvested by centrifugation at 7500 rpm for 10 min. The bacterial pellet was resuspended in lysis buffer (25 mM Tris pH 8, 100 mM NaCl) and sonicated for 30 s, 60 % amplitude, 8 cycles. The resulting cell lysate was centrifuged at 15,000 g for 30 mins at 4 °C. The clarified medium was filtered through a 0.4 μm filter prior to loading on a GSTrap FF column (Cytiva). Protein was eluted with 20 mM reduced glutathione and injected onto a Superdex 200 increase 10/300 GL column (Cytiva). Purified protein was stored at −80 °C.

### Production of single domain camelid nanobodies against Themis

Single domain camelid nanobodies (Nb) against Themis were raised by immunizing llamas with recombinant Themis and were selected for binding to the Themis^1-563^-Grb2^FL^ complex by ELISA and BLI to select complex-specific Nbs. The sequence of Nb256 was cloned in a pET15b bacterial expression vector in frame with an N-terminal pelB leader sequence to direct the protein to the bacterial periplasmic space, followed by an N-terminal His_6_ tag and a caspase-3 cleavage site. BL21 T7 Express *lysY*/*I^q^* (NEB) were transformed with pET15.Nb256 and protein expression was induced at OD_600_ 0.6-0.9 with 1 mM IPTG at 28 °C for 18-24 h. Cells were pelleted at 7500 rpm for 10 min at 4 °C and cell supernatant was discarded. Cell pellets were resuspended in 80 mL purification buffer (25 mM Tris pH 8, 100 mM NaCl). Cells were lysed by sonication and cellular debris and insoluble material was removed by centrifugation at 15,000 g for 30 min at 4 °C. The clarified medium was filtered through a 0.4 μm filter prior to chromatographic purification steps. Nb256 was captured via its His_6_ tag on a HisTrap FF column (Cytiva). The column was washed with purification buffer containing 20 mM imidazole, followed by elution with buffer containing 500 mM imidazole. Eluted Nb256 was concentrated and injected on an SD75 Increase 10/300 GL column (Cytiva) pre-equilibrated in purification buffer. Fractions containing Nb256 were pooled and stored at −80 °C. Protein purity was evaluated by SDS-PAGE analysis.

### ProMacrobody generation

Nb256 was converted to PMb256 and expressed and purified as described previously^28^. Briefly, the Nb256 sequence was cloned into the expression vector pBXNPH3M (Addgene, #110099). N-terminal of the Nb256 insert sequence, the plasmid expresses a pelB leader sequence followed by a His_10_ tag, maltose binding protein (MBP) and 3C protease site. A truncated MBP (starting at Leu7) is encoded C-terminally to the Nb256 insert, fused by a rigid double proline linker. The resulting sequence of the linker region is VTVPPLVI, where VTV is the conserved Nb C-terminus, PP denotes the rigid proline linker and LVI is the truncated N-terminus of MBP.

PMb256 was expressed in MC1061 *E.coli* cells and induced with 0.2 % arabinose at OD_600_ >1. After 3.5 hours, cells were harvested and PMb256 was purified using the same protocol as described for Nb256. After IMAC elution, imidazole in the resulting protein fraction was removed using a HiPrep 26/10 Desalting column. The N-terminal His_10_ tag and N-terminal MBP was cleaved off by overnight digestion with HRV-3C protease (Thermo Fisher) at 4 °C. The N-terminal MBP moiety, undigested protein and the His-tagged enzyme were subsequently removed by IMAC. The flowthrough from the second IMAC step, containing the cleaved tagless protein was concentrated and injected onto a Superdex 200 increase 10/300 GL column and fractions containing the proteins of interest were pooled and stored at −80°C. The purity of all recombinant proteins was evaluated by SDS-PAGE analysis.

### Size exclusion chromatography coupled to multi angle laser light scattering (SEC-MALLS)

120 μg protein (80 μL at 1.5 mg mL^-1^) was injected onto a Superdex 200 increase 10/300 GL column pre-equilibrated with buffer (25 mM Tris pH 8.5, 100 mM NaCl, 2 mM DTT), connected to an Agilent HPLC system with an autosampler. The column was coupled to an ultraviolet detector (Agilent), a DAWN8 MALLS detector and an Optilab refractometer (Wyatt). Data was analysed using the ASTRA8.2 software (Wyatt). Recombinant chitinaseA from *Clostridium perfringens* (1 mg mL^-1^)was used as a standard to correct for band broadening^45^. ChitinaseA was used as a standard instead of bovine serum albumin (BSA) as it lacks disulfide linkages and resists reduction by DTT.

### Cryo-EM sample preparation data collection

The Themis^1-563^-Grb2^FL^ complex was incubated with a 2 molar excess of PMb256. The Themis^1-563^-Grb2-PMb256 complex was purified by SEC on a Superdex 200 Increase 10/300 GL column in buffer containing maltose to stabilize the MBP moiety (25 mM Tris pH 8, 100 mM NaCl, 2 mM DTT, 2 mM maltose). A 4 μL sample of purified complex was applied to a glow discharged R2/1 300 mesh holey carbon copper grid (Quantifoil Micro Tools GmbH) at a concentration of 2.6 mg mL^-1^. Immediately before cryo-EM grid preparation, 8 mM CHAPSO (Anatrace Inc.) was added to the sample to promote spreading of the sample on the grid and reduce particle absorption at the air-water interface^46^. For the Themis^1-563^-PMb256 complex, a 4 μL sample of purified complex was applied to a glow discharged R2/1 300 mesh holey carbon copper grid (Quantifoil Micro Tools GmbH) at a concentration of 4.1 mg mL^-1^ in the presence of 8 mM CHAPSO, as before. Grids were blotted for 5 s and plunge frozen in liquid ethane using a Leica EM GP2 Plunge Freezer operated at 95 % humidity at room temperature.

Grids were screened using a JEOL 1400 Plus (VIB Bioimaging Core, Ghent and BECM, Brussels). Data collection was performed at the VIB-VUB Biological Electron Cryogenic Microscopy facility on a 300 kV CryoARM300 microscope (JEOL) equipped with an Omega filter (20 eV slit, JEOL) and K3 direct electron detector (Gatan) using SerialEM v3.8.17. Sixty frame movies were collected with a total exposure dose of 61.8 e^-^ A^-2^, at a magnification of 60,000, corresponding to a pixel size of 0.755 Å/pixel. For the Themis^1-563^-Grb2-PMb256 complex a total of 27,698 movies were collected, while for the Themis^1-563^-PMb256 complex, 7,755 movies were collected.

### Cryo-EM image processing

Processing of recorded movies was performed in cryoSPARC v4.5.1^47,48^. Movies were motion corrected and dose weighted using patch motion correction, and contrast transfer function (CTF) estimation on the aligned and dose-weighted micrographs was performed using patch CTF estimation. For all datasets, particle picking was performed using crYOLO^49^. The crYOLO-picked particle set was imported in cryoSPARC v4.5.1 and extracted using a box size of 416 pixels, downsampled to a box size of 208 pixels, corresponding to a pixel size of 1.51 Å per pixel (2x binned). The extracted particle stack was trimmed using several rounds of iterative 2D classification and 2D class selection to remove noisy particles. The cleaned particle stack was then used to train a TOPAZ picking model^50^ to identify additional particles. Particles extracted from TOPAZ and crYOLO picking were combined, duplicates were removed and more rounds of iterative 2D classification and class selection were performed. The resulting particle stack was used for *Ab-Initio* model generation using 3 classes as input, followed by heterogenous refinement of the resulting *Ab-Initio* volumes.

For the Themis^1-563^-Grb2^FL^-PMb256 complex, the dominant class from *Ab-Initio* reconstruction clearly corresponded to the complex, as further refinement of the other classes yielded very low resolution, noisy volumes. The class corresponding to the complex was further refined using non-uniform (NU) refinement, resulting in a map with a resolution of 3.3 Å, based on the 0.143 gold-standard Fourier shell correlation (FSC) criterion^51^. The particles contributing to this map were reextracted in a box size of 416 pixels corresponding to a pixel size of 0.755 Å per pixel and used for a final NU 3D refinement, resulting in a final map with a resolution of 3.3 Å based on the 0.143 gold-standard FSC criterion. Local refinement was performed by employing a soft mask around Themis^1-563^-Grb2^SH3C^. During local refinement, the rotation and shift search extent were set to 5° and 5 Å respectively, and a pose/shift gaussian prior was enabled with standard deviations of priors over rotation/shifts set to 15° and 7 Å. Local refinement resulted in a 3D reconstruction with a resolution of 3.2 Å and improved resolution at the Themis^1-563^-Grb2^SH3C^ interface. A comprehensive overview of the data processing workflow can be found in Extended Data Figure 3.

To investigate the conformational heterogeneity of the Grb2^SH3N^ and Grb2^SH2^ domains, particles contributing to the consensus 3D reconstruction were further analyzed using 3D classification without alignment using 10 classes as input. In the resulting volumes, 4 classes exhibited clear density corresponding to the N-terminal domains of Grb2. A predicted model of the Themis^1-563^-Grb2^FL^ complex was generated using Alphafold3 (AF3)^29^ and used to approximately fit the Grb2^SH3N^ and Grb2^SH2^ to each of the resulting volumes from 3D classification to make a soft mask in UCSF ChimeraX^52^. Particle subtraction was used to subtract the PMb256, which also displayed conformational heterogeneity, and local refinement using the Themis^1-563^-Grb2^FL^ soft masks was performed to increase the resolution of Grb2^SH3N^ and Grb2^SH2^ domains. NU refinement of each class resulted in maps with resolutions ranging between 3.8 Å to 4.3 Å. The 3D classification workflow can be found in Extended Data Fig. 9.

For the Themis^1-563^-PMb256 complex, the combined crYOLO- and TOPAZ-picked particles were used as input for *Ab-Initio* model generation using 5 classes as input, followed by heterogenous refinement of the resulting ab-initio volumes. The dominant class from *Ab-Initio* reconstruction was further refined using NU refinement, resulting in a map with a resolution of 5.8 Å, based on the 0.143 gold-standard FSC. Particles were further analysed using 3D classification without alignment followed by heterogeneous refinement, resulting in two dominant classes. NU refinement of each class resulted in maps with a resolution of 6.3 Å for class 1 (CABIT1 domain present) and 7 Å for class 2 (CABIT1 domain absent). Particles from class 2 were subjected to 3D variability analysis using 3 orthogonal principle modes, filtered to 10 Å resolution. An overview of the data processing workflow can be found in Extended Data Figure 8.

All final maps were post-processed using DeepEMhancer^53^ for the purpose of model building and visualization.

### Cryo-EM model building and refinement

A structural model of the Themis^1-563^-Grb2^FL^ complex was predicted using Alphafold-Multimer version 2.1^54^. A predicted model of Nb256 was made with ESMFold^55^, along with the previously published PMb structure (PDB 7omt)^28^. These predicted and experimental models were combined to make an initial model for the Themis^1-563^-Grb2-PMb256. This initial model was rigid-body fitted in the sharpened map using UCSF ChimeraX and subsequently subjected to automatic molecular dynamics (MD) flexible fitting using NAMDINATOR^56^. The resulting flexibly fitted complex was then used for several rounds of manual building in Coot^57^ using the DeepEMhancer sharpened map to guide model building. The manually built model was then subjected to several cycles of real-space refinement in Phenix^58^ (version 1.21.1-5286) using the regularly sharpened map, enabling global minimization, local grid search, atomic displacement parameter (ADP) refinement, secondary structure and Ramachandran restraints, and using a nonbonded weight parameter of 300. Validation of the model was performed using Phenix and MolProbity^59^. Cryo-EM data collection, refinement and validation statistics are summerised in Table 1. Visualization of cryo-EM maps and structural models was performed using UCSF ChimeraX and Pymol (The PyMOL Molecular Graphics System, version 2.5.8, Schrödinger, LLC).

### In silico structure prediction and bioinformatic analysis

A structural model for the complex of Themis^1-563^-Grb2^FL^ was predicted using Alphafold-Multimer version 2.2^54^ and used to generate a starting model for real-space cryo-EM refinement. A structural model for Nb256 was predicted using ESMFold^55^.

Multiple sequence alignment was performed using Clustal Omega^60^ and visualized using ESPript3.0^61^. PDBePISA^62^ was used for protein-protein interaction analysis, and interfacing residues were indicated on the sequence alignment.

### Isothermal Titration Calorimetry (ITC)

Proteins used in ITC experiments were expressed in HEK293S suspension cells, as previously described. As a final purification step, all proteins were buffer exchanged to the same buffer (25 mM Tris pH8, 100 mM NaCl, 2 mM Tris(2-carboxyethyl)phosphine (TCEP)). TCEP is used instead of DTT as a reducing agent here as DTT has been suggested to cause baseline shifts and background signals in ITC. Experiments were performed using a MicroCal PEAQ-ITC instrument at 23 °C. Titrations were preceded by an initial injection of 0.4 μL. Titrations consisted of 12 injections of 3 μL, with 150 s spacing between injections. Throughout the titration the sample was stirred at a speed of 750 rpm. Data was analyzed using PEAQ-ITC analysis software (version 1.1.0.1262, Malvern) and fit using a ‘one set of sites’ model.

### Biolayer interferometry (BLI)

Characterization of the Themis^1–563^-Grb2 interaction and screening of mutant Themis and Grb2 variants was performed by immobilizing biotinylated wild-type (WT) or mutant Grb2 variants on streptavidin (SA) biosensors (Sartorius). To this end, WT Grb2 and mutant variants (Y160A, F165A, R179A, F182A, W193A, Y2909A) were generated by overlap extension PCR^63^ and cloned in a modified pHLSec vector, lacking a secretion signal, in frame with an N-terminal Avi-tag. All constructs were transiently co-transfected in suspension-adapted HEK293S cells together with a BirA expression plasmid^64^ and supplemented with 50 μM biotin upon transfection to specifically biotinylate the Avi-tag. After 2 days, cells were harvested and biotinylated Grb2 variants were purified by IMAC and SEC, as previously described. All mutant variants were stable and had comparable yields and biochemical behavior as WT.

All measurements of binding kinetics and dissociation constants were performed using an Octet Red 96 (Sartorius) in kinetics buffer (25 mM Tris pH8, 100 mM NaCl, 1 % (w/v) BSA, 0.02% (v/v) Tween 20). SA biosensors were functionalized with biotinylated Grb2 WT or mutant variants to an optical shift of 1 nm. To exclude any non-specific binding, all experiments were performed with double referencing, where non-functionalized biosensors were used as a control by measuring in parallel all ligand concentrations and kinetics buffer alone. All data was fitted using the Forté Bio Data Analysis Software 9.0 (Sartorius), utilizing a 1:1 interaction model.

Interaction studies with PRS peptides from Themis and Sos1 were carried out with biotinylated peptides immobilized on SA biosensors, as before. Sos SH3 domain inhibitor peptide (Santa Cruz) was *in vitro* biotinylated at its amino terminus using EZ-Link NHS-PEG4-Biotin (Thermo Fisher). Excess biotin was removed by injection on an Enrich70 column (Biorad). N- terminally biotinylated Themis^PRS^ peptide was purchased from GenScript.

For measurements of binding kinetics and dissociation constants involving the isolated Grb2^SH3C^ domain, GST-tagged Grb2^SH3C^ was immobilized on GST biosensors (Sartorius) and measured as above.

### CRISPR-mediated gene deletion

sgRNAs against Themis were designed using CHOPCHOP^65^ and cloned into pSpCas9(BB)- 2A-GFP vector (Belgian Coordinated Collections of Microorganisms, BCCM). The sgRNA oligos used were as follows:

ligo 1: 5’- CACCG GGATGACAGCTTGTCATGAG – 3’
ligo 2: 5’- AAAC CTCATGACAAGCTGTCATCC C – 3’

Jurkat E6.1 cells were cultured in RPMI 1640 medium (Gibco) supplemented with 10 % fetal calf serum (FCS, Biowest). Jurkat cells were electroporated with 2 μg of plasmid containing sgRNA with Amaxa 2D nucleofector using Cell Line Nucleofector Kit V (Lonza Bioscience). After 72 h, transfected cells were sorted for GFP positivity using an FACS ARIAII cell sorter (BD Biosciences). Sorted cells were further cultured as a bulk population. In parallel, cells were also transfected with an empty vector without sgRNA as a control cell line. Gene knockout was confirmed at the protein level by western blot analysis. All cells were cultured at 37 °C in a humidified atmosphere with 5 % CO_2_.

### Lentiviral reconstitution of Themis mutants

To investigate the role of Themis in T cell receptor signalling, Themis WT or mutant variants (I181A, F327A, E483A, Y541A, R544A, R545A, P554A, P557A, I181A/R545A, P554A/P557A) were cloned in the pRRLSIN.cPPT.PGK-GFP-T2A.WPRE lentiviral vector. Lentiviral production was carried out in the Lenti-X™ 293T cell line (Takara) using TransIt-Lenti transfection reagent (Mirus Bio) at a ratio of 1 μL TransIt-Lenti to 2 μg DNA. Briefly, 2 μg DNA (0.9 μg packaging vector, 0.1 μg envelope vector, 1 μg lentiviral vector containing Themis variants or empty vector) was diluted in OptiMEM (Thermo Fisher) to a final volume of 200 μL. The TransIt-Lenti reagent was added to the DNA and incubated for 10 min at room temperature. The TransIt-Lenti/DNA mix was added drop-wise to cells. After 48 hours, cell culture medium containing viral particles was harvested, centrifuged to remove cellular debris and stored at −80 °C.

The Themis-deficient Jurkat cell line generated by CRISPR editing was transduced with the lentiviral particles containing WT or mutant variants of Themis. Themis-deficient Jurkat cells were seeded at 200,000 cells per well in a 24 well plate coated with retronectin (Takara) and 400 μL of lentiviral supernatant was added. After 48 hours, transduction efficiency (GFP positive cells) was checked by flow cytometry. Transduced cells were further cultured and expanded and after 7 days, transduced cells were sorted for similar levels of GFP positivity using a BD FACS Melody (BD Biosciences). Sorted cells were further cultured as a bulk population. In parallel, Jurkat CRISPR control cells were also transduced with an empty pRRLSIN.cPPT.PGK-GFP-T2A.WPRE vector. Reconstituted Themis expression was confirmed at the protein level by western blot analysis. All cells were cultured at 37 °C in a humidified atmosphere with 5 % CO_2_.

### T cell activation assays

Jurkat cells (1 x 10^6^) were stimulated with 2 μg/mL plate-bound anti-CD3 antibody (BD Biosciences) and 1 μg/mL soluble anti-CD28 antibody (BD Biosciences) for 30-60 s. At the indicated timepoints, cells were lysed with 100 μL 2X SDS loading buffer and incubated on ice for 10 min. Samples were boiled at 95 °C and 15 mL was loaded on Criterion TGX Precast mini protein gels (Biorad). Gels were transferred on nitrocellulose membranes using the Trans-Blot turbo transfer system (Biorad). Membranes were blocked for 1 hour in 5% skimmed milk powder in Tris-buffered saline containing 0.05 % Tween 20 (TBST). The indicated proteins were probed using specific primary antibodies, typically diluted 1:1000, overnight at 4 °C. Membranes were washed for three times with TBST and incubated with the relevant horseradish peroxidase-conjugated secondary antibody. Membranes were washed and proteins were visualized using ECL Prime western blotting system (Cytiva) on an Amersham imager 600 (GE Healthcare Life Sciences).

The primary antibodies used are as follows: anti-Themis rabbit monoclonal (catalog number ab126771, Abcam), pSHP-1 (Tyr564, clone D11G5, catalog number 8849, Cell Signalling Technology), SHP-1 (clone C14H6, catalog number 3759, Cell Signalling Technology), pERK (Thr202/Tyr204 p-p44/p-p42 MAPK, clone D13.14.4E, catalog number 4370, Cell Signalling Technology), ERK (p44/p42 MAPK, catalog number 9102, Cell Signalling Technology), β- tubulin (ab21058, Abcam). The secondary antibody used was Peroxidase AffiniPure goat anti-rabbit IgG (Jackson Immunoresearch).

### Animals

Themis^-/-^ and Themis^L411V^ transgenic mice were generated by CRISPR-Cas9 editing by the VIB Transgenic Core Facility (Ghent, Belgium). All mice and their littermate controls were on a C57BL/6J background. All transgenic mice were bred in-house at the VIB Centre for Inflammation Research in a specific pathogen-free (SPF) environment. All mouse experiments were conducted according to institutional, national and European animal regulations. All experimental animal procedures were approved by the Ethics Committee of Ghent University (EC22-61).

### Immunophenotyping by flow cytometry

Thymic lobes and spleens from age-matched adult mice (8-12 weeks) were surgically removed and processed into single-cell suspensions by pressing the organs through 70 μm cell strainers (VWR International) in phosphate-buffered saline (PBS) supplemented with 2 % FCS. Spleens were further processed with ACK lysis buffer (Westburg b.v.) for 3 min at room temperature to remove red blood cells.

Subsequent flow cytometry analysis of all samples was performed on ice. Antibody staining was performed for 30 min in the dark. Fixable Viability Dye eFluor 780 (Thermo Fisher) was used to exclude dead cells in all flow cytometry experiments. Single cell suspensions were analysed on a FACSymphony A5 (BD Biosciences). The antibodies used for flow cytometry were as follows:

| Antigen | Fluorochrome | Clone | Source | Catalog no. | Dilution |
| --- | --- | --- | --- | --- | --- |
| CD117 | BB515 | 2B8 | BD Biosciences | 564481 | 1:100 |
| CD44 | BV421 | IM7 | BD Horizon | 560451 | 1:400 |
| CD8β | BV510 | YTS156.7.7 | Biolegend | 126631 | 1:400 |
| CD25 | BV605 | PC61 | BD Horizon | 563061 | 1:200 |
| TCRβ | BV711 | H57-597 | BD Horizon | 563135 | 1:100 |
| CD11c | BV785 | N418 | Biolegend | 117336 | 1:400 |
| CD19 | BV785 | B4 | Biolegend | 115543 | 1:200 |
| CD11b | BV785 | M1/70 | Biolegend | 101243 | 1:400 |
| CD4 | AF647 | GK1.5 | Biolegend | 100424 | 1:400 |
| CD62L | FITC | MEL-14 | Thermo Fisher | 11-0621-82 | 1:400 |
| NK1.1 | PE-Cy7 | PK136 | BD Pharmingen | 552878 | 1:100 |
| CD3 | BUV395 | 145-2C11 | BD Horizon | 563565 | 1:50 |
| CD24 | BUV737 | M1/69 | BD Horizon | 565308 | 1:200 |
| CD27 | BUV737 | LG.3A10 | BD Horizon | 565307 | 1:200 |

### Induction and assessment of arthritis

For the induction of collagen-induced arthritis (CIA), male mice were immunized intradermally at the base of the tail with 200 μg of chicken type II collagen (MD Biosciences) dissolved in 0.1 M acetic acid emulsified in complete Freund’s adjuvant (BD Biosciences). On day 21 postimmunization, mice received a second challenge using the same mixture. Thereafter, mice were monitored by 2 independent investigators for clinical symptoms of arthritis until day 60, at which time they were euthanized. Three joints were evaluated for clinical scoring of arthritis, namely the interphalangeal joints (digits), metacarpophalangeal joints and carpal/tarsal joints. Clinical symptoms were scored on a scale of 0-3, in which 0 = normal, 0.5 = swelling and redness of 1 interphalangeal joint, 1 = swelling and redness of at least 2 interphalangeal joint or the metacarpophalangeal joint or carpal/tarsal joint, 2 = swelling and redness of 2 joints, 3 = swelling and redness of the entire paw.

For histopathologic evaluation of inflammation, mouse knees were fixed in 4 % formaldehyde and decalcified in 5 % formic acid. Paraffin-embedded sections were stained with hematoxylin and eosin and assessed for severity of inflammation and joint damage. To measure serum anti-type II collagen antibody levels, blood samples were collected at day 60 and allowed to clot at room temperature. After clotting, samples were centrifuged and stored at −20 °C. Anti-type II collagen antibodies were measured by ELISA by coating 96-well half area microplates (Greiner Bio One) with chicken type II collagen (1.26 μg mL^-1^) for 2 hours at 37 °C. Non-specific binding was blocked with 0.1 % casein in PBS for 1 hour at 37 °C. Samples were diluted 1:10 and incubated at 4 °C, overnight. Detection antibodies coupled to HRP (IgG1-HRP and IgG2α-HRP, Southern Biotech) were added for 1 hour at room temperature. Finally, TMB substrate solution (OptEIA TMB substrate reagent set, BD Biosciences) was added to the wells and the reaction was stopped with 1 M H_2_SO_4_. OD values were measured at 450 nm.

### Data availability

Cryo-EM maps have been deposited in the Electron Microscopy Data Bank (EMDB) and the Protein Data Bank (PDB) under accession codes: PDB 9i3p, EMD-52603 (Themis^1-563^-Grb2-PMb256 complex), PDB 9iaz, EMD-52787 (Themis^1-563^-Grb2-PMb256 complex, local refinement), EMD-52604 (Themis^1-563^-Grb2-PMb256 complex, 3D classification Class 1), EMD-52605 (Themis^1-563^-Grb2-PMb256 complex, 3D classification Class 2), EMD-52606 (Themis^1-563^-Grb2-PMb256 complex, 3D classification Class 3), EMD-52607 (Themis^1-563^- Grb2-PMb256 complex, 3D classification Class 4), EMD-52608 (unbound Themis^1-563^- PMb256 complex), EMD-52609 (unbound Themis^1-563^- PMb256 complex, 3D classification Class 1), EMD-52609 (unbound Themis^1-563^- PMb256 complex, 3D classification Class 2).

## AUTHOR CONTRIBUTIONS

D.M.C. performed recombinant protein production, biophysical characterization by SEC-MALLS, BLI and ITC, cellular assays, structural studies by cryo-EM and data analysis. A.S. assisted with recombinant protein production. E.G. performed the rheumatoid arthritis mouse model experiments, with supervision by D.E. P.T performed immunophenotyping experiments, with supervision by P.V. S.S and J.B. provided critical reagents and expertise. I.V. and Y.V.D. provided reagents and performed viral transductions, with expertise and supervision from T.T.J.F. and Y.B. assisted with cryo-grid preparation, cryo-EM data collection and analysis. R.M. and A.F.C. assisted with initial construct design and protein purification. D.M.C. and S.N.S. wrote the manuscript with contributions from all authors. S.N.S. conceived and supervised the project.

## ACKNOWLEDGEMENTS

We thank M. Fislage at the VIB-VUB facility for Biological Electron Cryogenic Microscopy for assistance with cryo-EM data collection, technical support and infrastructural access. We thank K. Balcaen and J. Haustraete at the VIB Protein Core Facility for access to the SEC-MALLS instrument and technical support. We are grateful to the staff at VIB Bioimaging Core for access to the cryo-EM screening microscope and VIB Flow Core Ghent for technical support. We thank K. Staes and T. Hochepied from the VIB Transgenic Mouse Core for generating the Themis KO and ThemisL411V transgenic mouse lines. We thank S. Schoonooghe and R. Hassanzadeh for generating the nanobody library and providing technical assistance. The pGEX plasmid containing Grb2^SH3C^ was a kind gift from Prof. Stephan M. Feller (Martin Luther University, Halle Wittenberg). D.M.C. is a Research Foundation Flanders senior postdoctoral research fellow (FWO grant number 1219923N). S.N.S. acknowledges support from Research Foundation Flanders (FWO grants G049820N, G0H1222N) and the Flanders Institute for Biotechnology (VIB grant C0101VIB).

## EXTENDED DATA FIGURES

**Extended Data Fig. 1.**
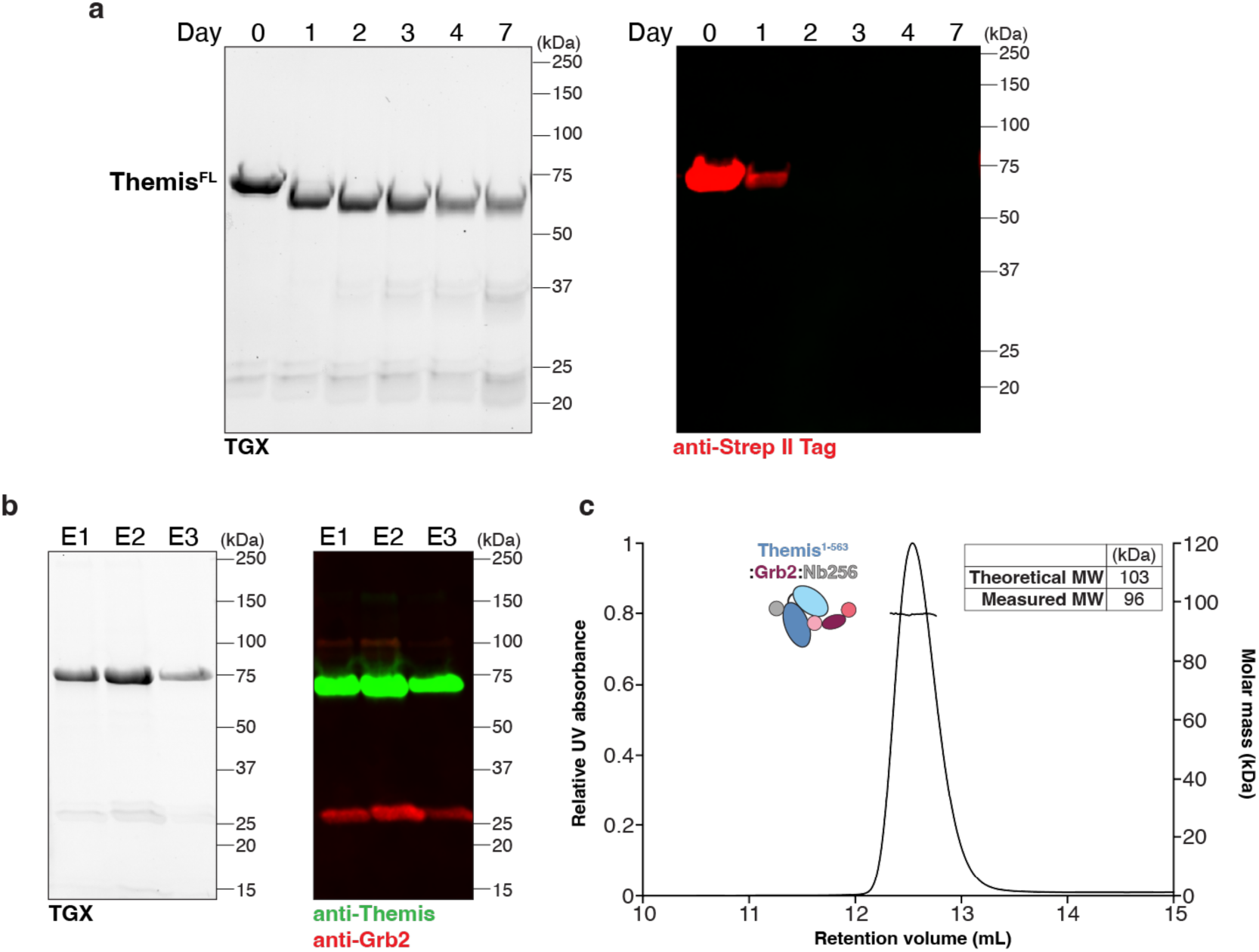
Recombinant Themis co-purifies with endogenous Grb2. **a,** Full-length C-terminal StrepII-tagged Themis (Themis^FL^) was expressed and purified from HEK suspension cells. Purified recombinant Themis^FL^ was incubated at room temperature and samples were taken at indicated timepoints to check protein stability. Representative TGX gel and western blot analysis with an anti-StrepII tag antibody is shown. The predominant band at approximately 73 kDa corresponds to purified Themis^FL^. Themis^FL^ loses its C-terminal StrepII tag and moves to a lower molecular weight (MW) over time. **b,** Themis was expressed and purified from HEK suspension cells. Representative TGX gel analysis of elution (E) fractions after affinity purification. The same gel was probed with specific anti-Themis and anti-Grb2 antibodies and detected with fluorescently tagged secondary antibodies. A fraction of endogenous cellular Grb2 was pulled down during Themis^FL^ purification from HEK suspension cells. **b,** SEC-MALLS analysis of co-expressed and purified Themis^1-563^-Grb2 in complex with Nb256. The Themis^1-563^-Grb2-Nb256 complex has a theoretical mass of 103 kDa.

**Extended Data Fig. 2.**
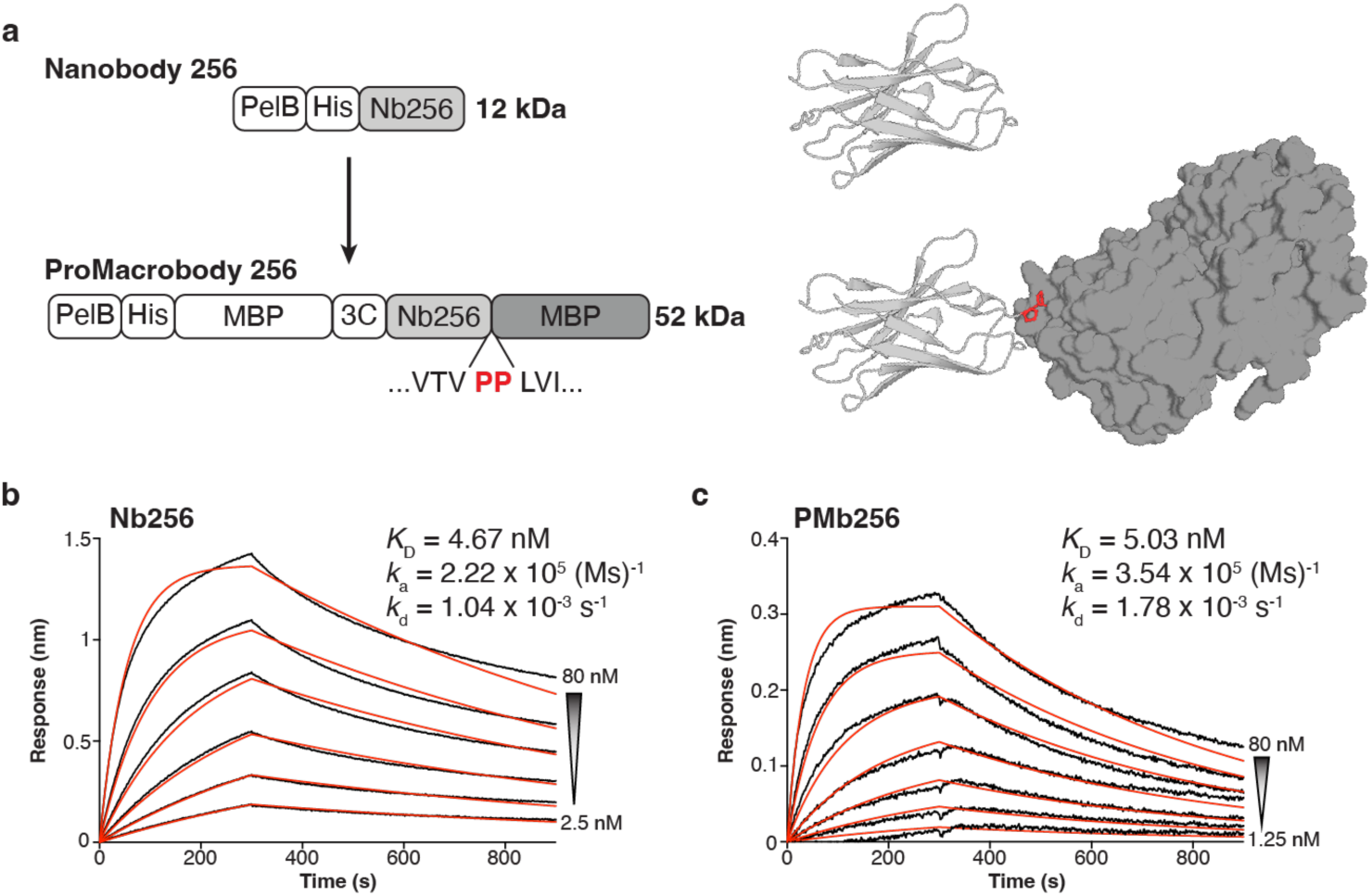
Conversion of Nb256 to PMb256. **a,** Comparison of the construct layout for the expression of Nb256 and PMb256. The original Nb construct contained a pelB leader sequence for periplasmic expression, followed by a hexahistidine (His6) tag and the Nb256 sequence. To generate PMb256, the Nb256 sequence was cloned into a pBXNPH3M expression vector containing a PelB leader sequence, followed by a decahistidine (His10) tag, MBP and 3C protease site. A truncated MBP moiety starting at residue Leu7 (LVI…) is encoded C-terminal to the Nb256 sequence, connected by a rigid double proline linker. Cartoon representations of Nb256 and PMb256 are shown. Structures were extracted from the cryo-EM model of the Themis^1-5^^63^-Grb2-PMb256 complex. The double proline linker is coloured in red and the MBP moiety is shown in surface representation. **b,** Representative BLI response curve to characterize binding of Themis1-563 to immobilized Nb256. **c,** Representative BLI response curve to characterize binding of Themis1-563 to immobilized PMb256. Response curves were fitted with a 1:1 model (red) to quantify the kinetics (*k*_a_, *k*_d_) and binding affinity (*K*_D_).

**Extended Data Fig. 3.**
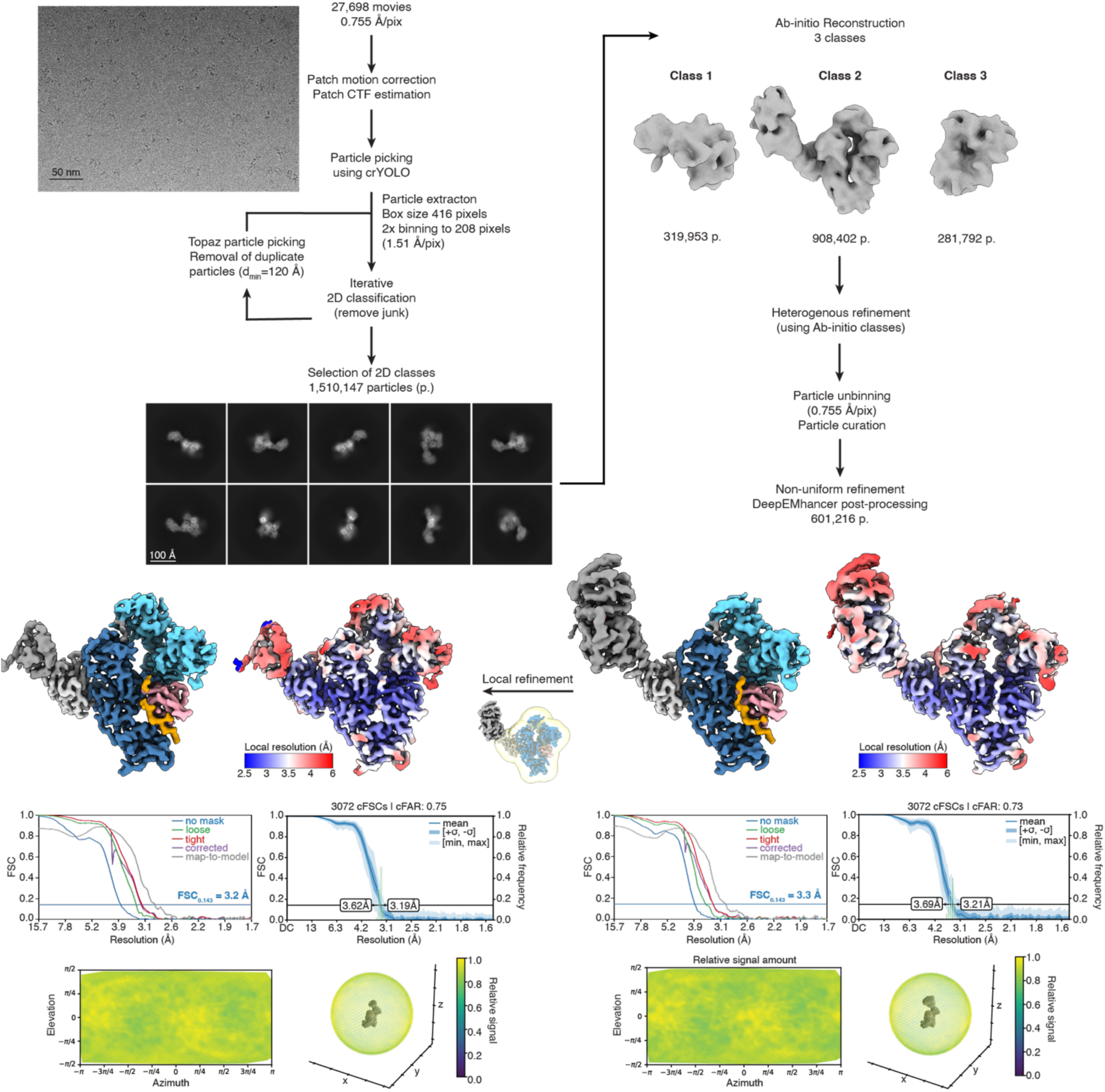
Cryo-EM data processing workflow for the Themis^1-563^-Grb2-PMb256 complex. Data processing was performed in CryoSPARC v4.5.1^47,48^ and map post-processing was performed using DeepEMhancer^53^. Gold standard Fourier Shell Correlation (FSC) curves are shown and the estimated resolution at FSC = 0.143 (blue line) is shown for the corrected FSC curves (purple line).

**Extended Data Fig. 4.**
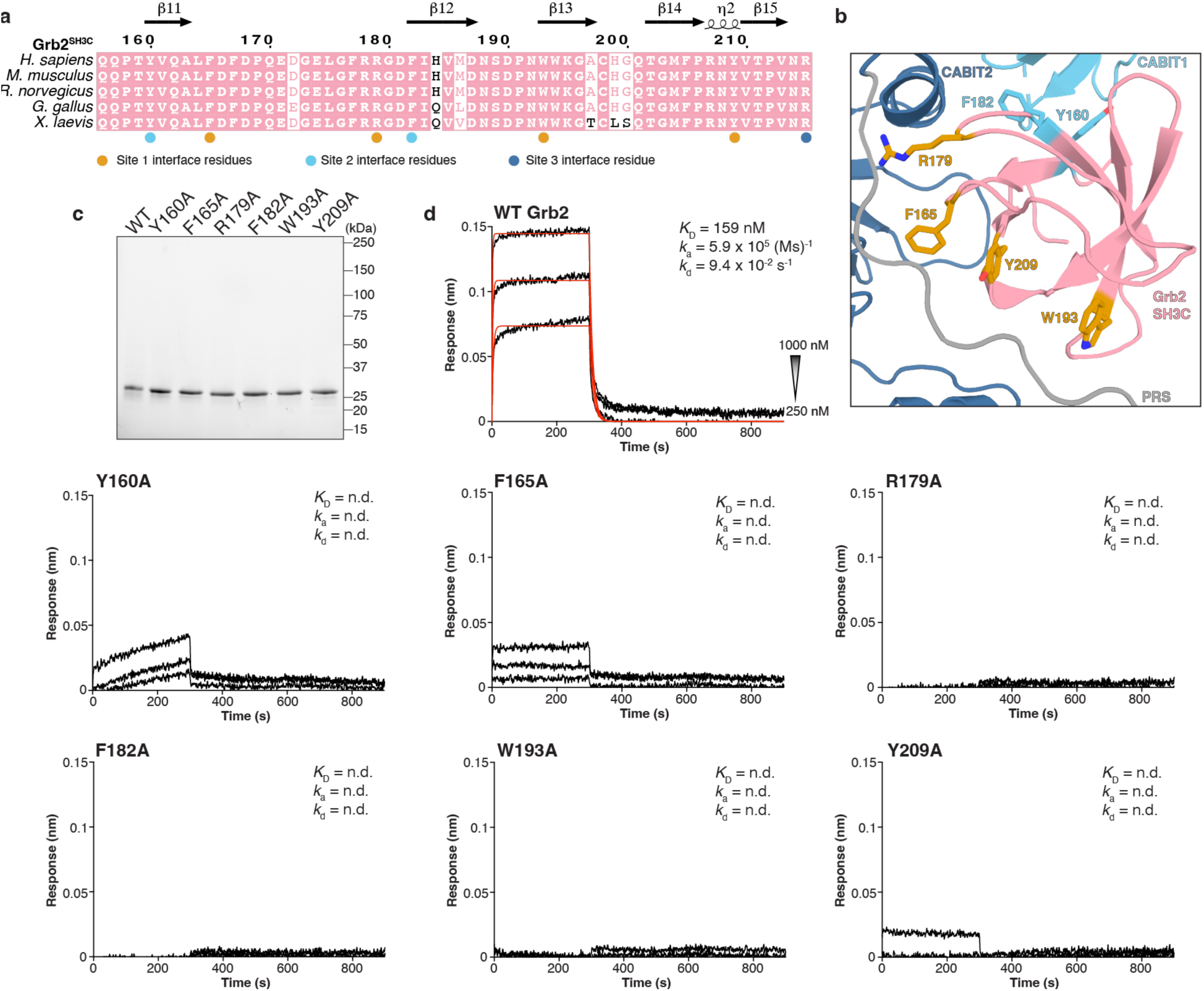
Functional interrogation of Grb2^SH3C^ mutants. **a,** Sequence alignment of various vertebrate Grb2^SH3C^ sequences using the ESPripT server (https://espript.ibcp.fr) and structural annotation according to secondary structure elements. Coloured symbols indicate residues participating in interaction sites 1-3. *H. sapiens* (Homo sapiens); *M. musculus* (Mus musculus); *R. norvegicus* (Rattus norvegius); *G. gallus* (Gallus gallus); *X. laevis* (Xenopus laevis). **b,** Zoom-in view of the Themis^1-563^-Grb2^SH3C^ interaction interfaces. Site 1 and site 2 interface residues targeted by mutagenesis are shown in stick representation and coloured as in **a**. Grb2^SH3C^, Themis^PRS^, Themis^CABIT1^, Themis^CABIT2^ are shown in pink, orange, cyan and dark blue, respectively. **c,** Representative SDS-PAGE analysis of purified Grb2^SH3C^ mutants used in BLI experiments. All mutant variants displayed similar biochemical behaviour to wild-type (WT) Grb2 **d,** BLI response curves for the interaction of immobilized WT or mutant Grb2FL variants with Themis^1-563^. A 1:1 binding model was fitted (in red) to quantify the kinetics (*k*_a_, *k*_d_) and binding affinity (*K*_D_). For all Grb2 mutants tested, kinetics and affinity could not be determined (n.d.) due to no binding or poor data fitting.

**Extended Data Fig. 5.**
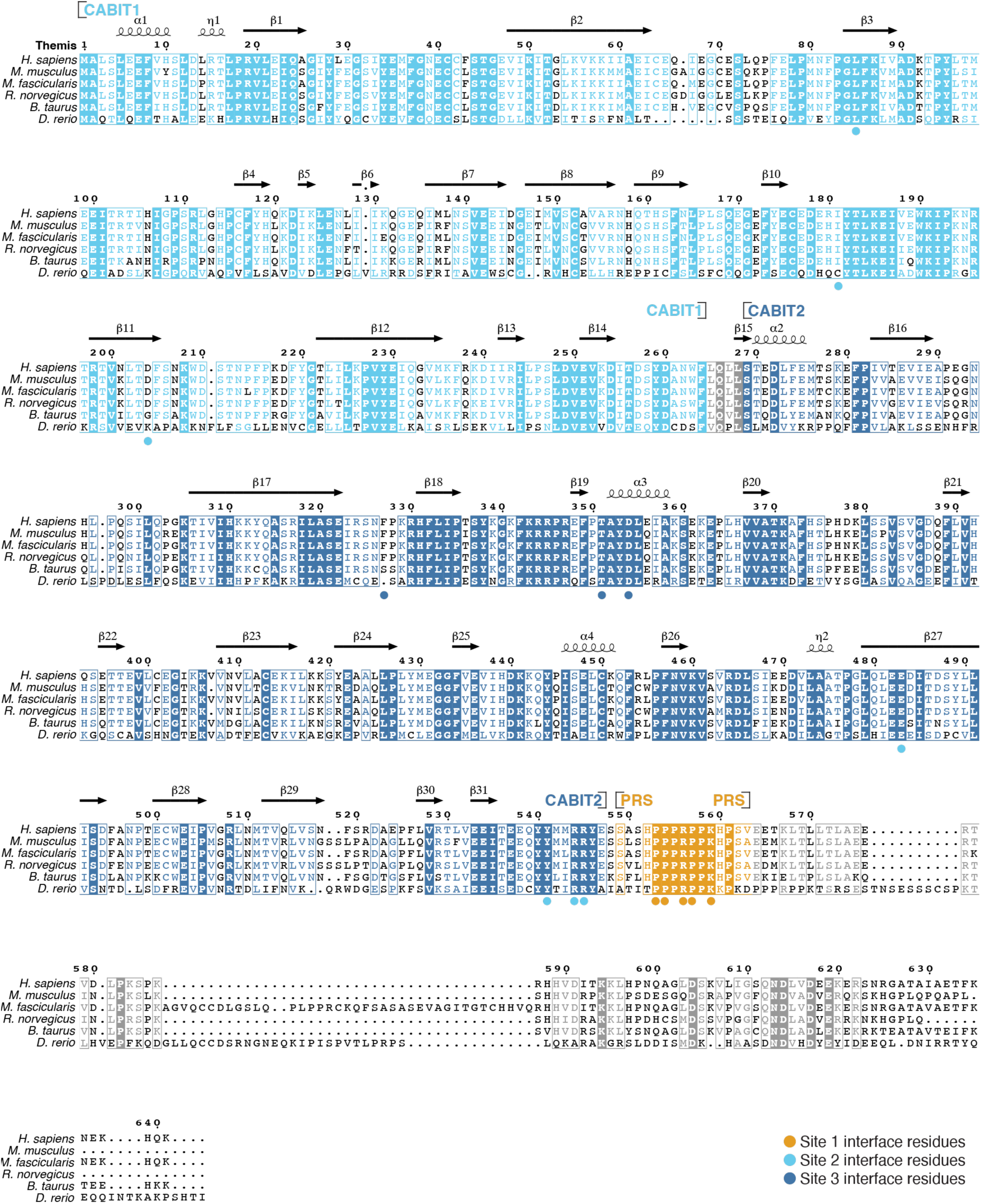
Conservation of Grb2^SH3C^ interfacing residues in Themis. Structurally annotated multiple sequence alignment of various vertebrate Themis sequences using the ESPripT server (https://espript.ibcp.fr). Coloured symbols indicate residues participating in interaction sites 1-3. CABIT1, CABIT2 and PRS domain delineations are indicated. *H. sapiens* (Homo sapiens); *M. musculus* (Mus musculus); *M. fascicularis* (Macaca fascicularis); *R. norvegicus* (Rattus norvegius); *B. taurus* (Bos taurus); *D. rerio* (Danio rerio).

**Extended Data Fig. 6.**
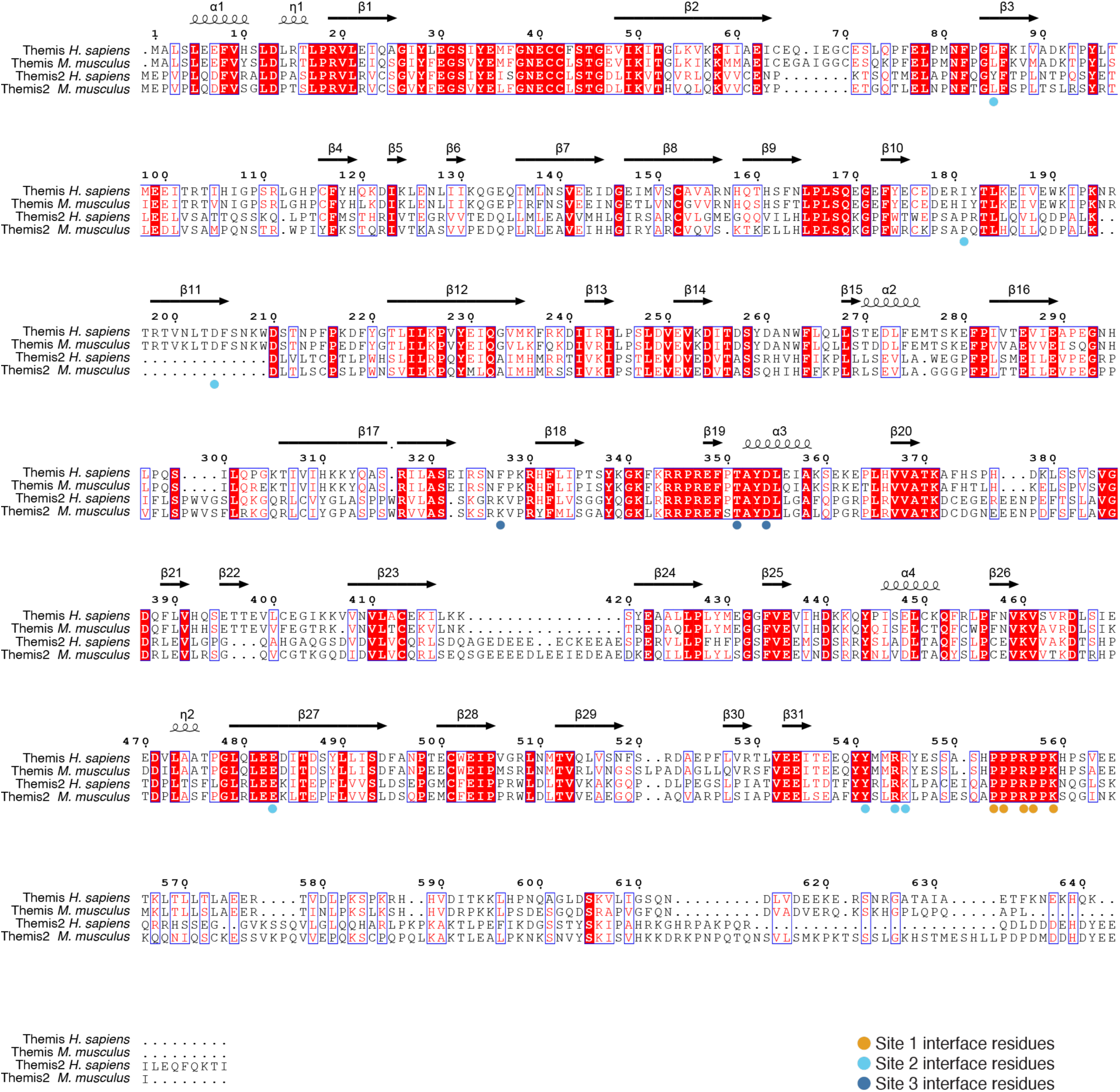
Sequence alignment of Themis and Themis2. Structurally annotated multiple sequence alignment of Themis and Themis2 sequences using the ESPripT server (https://espript.ibcp.fr). Coloured symbols indicate residues participating in interaction sites 1-3 with Grb2SH3C. *H. sapiens* (Homo sapiens); *M. musculus* (Mus musculus).

**Extended Data Fig. 7.**
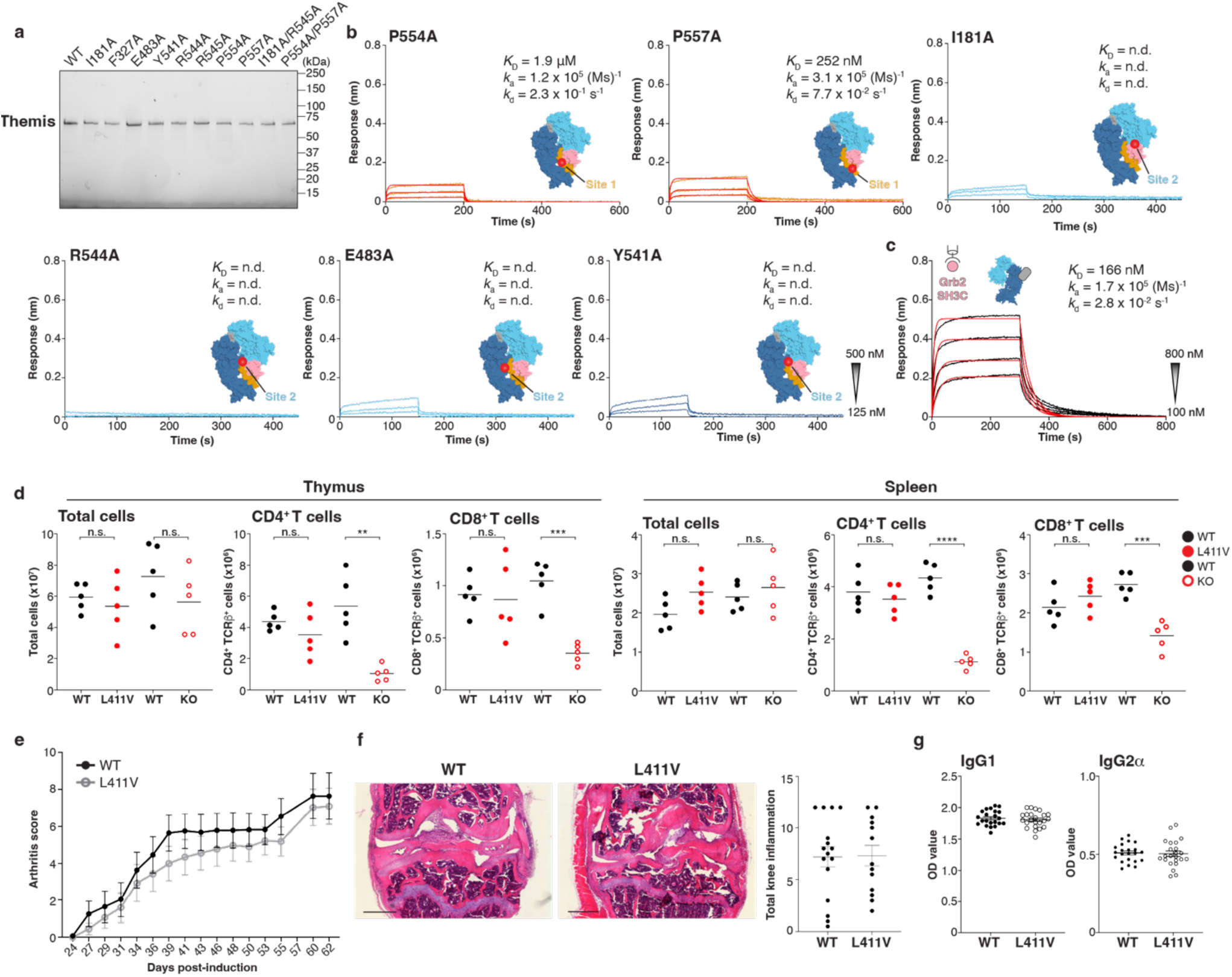
Functional interrogation of Themis mutants. **a,** SDS-PAGE analysis of purified Themis^1-563^ mutants used in BLI experiments. All mutant variants displayed similar biochemical behaviour to wild-type (WT) Themis^1-563^. **b,** BLI response curves for the interaction of immobilized WT Grb2^FL^ with Themis^1-563^ mutants. A 1:1 binding model was fitted (in red) to quantify the kinetics (*k*_a_, *k*_d_) and binding affinity (*K*_D_). Kinetics and affinity could not be determined (n.d.) for the indicated mutants due to no binding or poor data fitting. The location of each mutation in the Themis^1-563^-Grb2^SH3C^ interface are indicated on a surface representation of the complex. **c,** BLI response curve for the interaction of immobilized GST-Grb2^SH3C^ with Themis^1-563^ in complex with Nb256. A 1:1 binding model was fitted (in red) to quantify the kinetics (*k*_a_, *k*_d_) and binding affinity (*K*_D_). For all BLI experiments, start and end concentrations of the 2-fold dilution series is shown as an inset. **d,** Total cellularity and quantification of CD4^+^ and CD8^+^ single positive T cell populations in the thymus and spleen of Themis^KO^ and Themis^L411V^ mice compared to wild-type (WT) littermates by flow cytometry. Data are from one experiment with five mice per group. Bars represent the mean of the data. n.s. non-significant, **p < 0.01, ***p < 0.001, ****p < 0.0001 by two-tailed t-test. **e,** Clinical arthritis scores were assessed in the ankle joints of mice after induction of CIA, in groups of male Themis^L411V^ transgenic mice (n=23) and wild-type (WT) littermates (n=23). Results are the mean ± SEM. **f,** Knee joints of male WT or Themis^L411V^ transgenic mice were examined for histological features of arthritis to confirm the clinical data. Hematoxylin and eosin staining was carried out to assess the amount of infiltrate and exudate in the femorotibial joints. Scale bars represent 500 μm. **g,** Serum antibody levels of IgG1 and IgG2α anti-collagen antibodies from WT and Themis^L411V^ mice were measured by ELISA.

**Extended Data Fig. 8.**
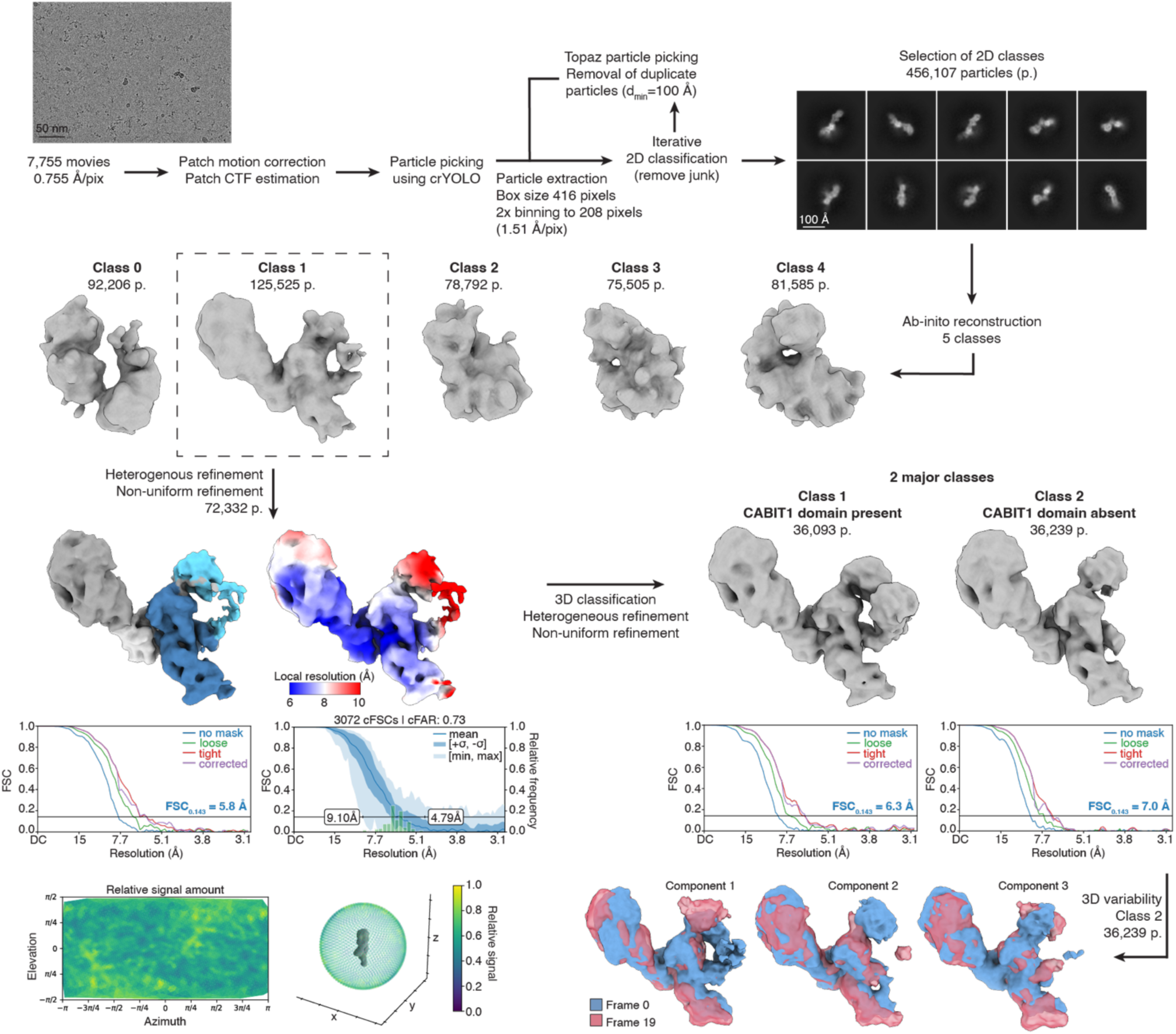
Cryo-EM data processing workflow for the Themis^1-563^-PMb256 complex. Data processing was performed in CryoSPARC v4.5.1. Sharpened maps from non-uniform refinement in CryoSPARC are shown. Gold standard Fourier Shell Correlation (FSC) curves are shown and the estimated resolution at FSC = 0.143 (blue line) is shown for the corrected FSC curves (purple line).

**Extended Data Fig. 9.**
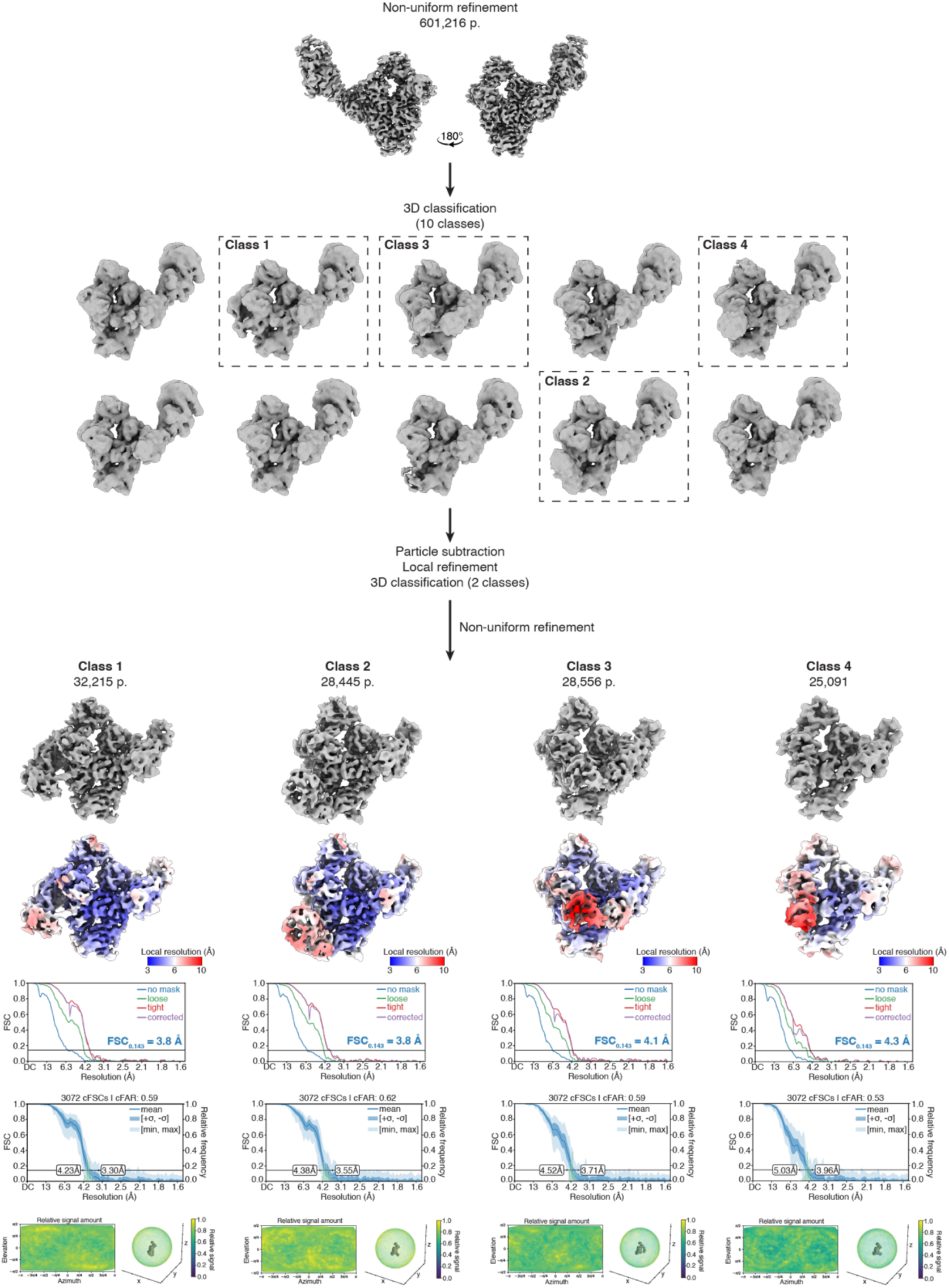
3D classification and data processing workflow to identify conformational heterogeneity of Grb2^SH3N^ and Grb2^SH2^ domains. Data processing was performed in CryoSPARC v4.5.1. Particles (601,216 p.) contributing to the Themis^1-563^-Grb2-PMb256 map were further analysed in a 3D classification job without alignment. The resulting 4 classes exhibiting volume corresponding to the N-terminal domains of Grb2 were subjected to particle subtraction to remove the MBP moiety from the PMb followed by local refinement. Final particle numbers (p.) and sharpened maps from non-uniform refinement for each class are shown. Gold standard Fourier Shell Correlation (FSC) curves are shown and the estimated resolution at FSC = 0.143 (blue line) is shown for the corrected FSC curves (purple line).

**Extended Data Fig. 10.**
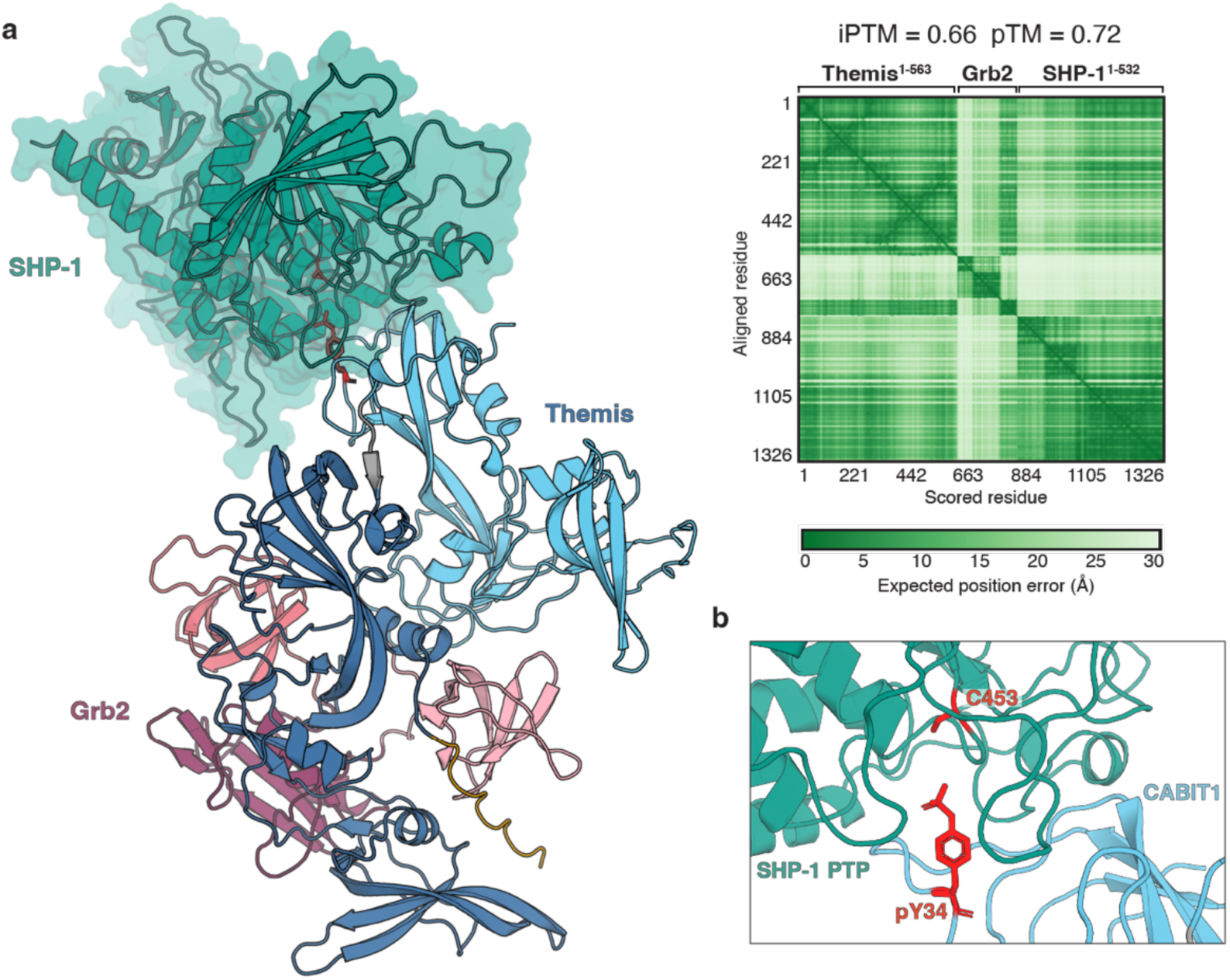
AlphaFold3 predicts the formation of a Themis-Grb-SHP-1 ternary complex. **a,** AlphaFold3 prediction of Themis^1-563^ in complex with Grb2^FL^ and SHP-1^1-5^^32^ using the Google Deepmind webserver (https://golgi.sandbox.google.com). Phosphorylation of Tyr34 (pY34) in Themis was included as a post-translational modification in the prediction for the complex. Truncated sequences for Themis and SHP-1 were used for prediction to exclude lower confidence scores caused by intrinsically disordered sequences at the C-termini of these proteins. Truncated SHP-1 lacking the flexible 61 residue C-terminal tail (residues 1-532) has previously been characterized using X-ray crystallography. Themis and Grb2^FL^ are coloured as before. SHP-1 is coloured in teal and depicted as a surface representation overlaying the cartoon representation. The iPTM and PTM scores and PAE plot are also included as confidence metrics. **b,** Zoom-in view of the predicted interface between Themis pY34 and the catalytic pocket of SHP-1 phosphatase (PTP) domain. Themis pY34 and the active site cysteine of SHP-1 (C453) are depicted in stick representation and coloured red.

**Extended Data Table 1.** Analysis of Themis^1-563^:Grb2:PMb256 interaction interfaces. Interacting residues in the Themis^1-563^:Grb2^SH3C^ and Themis^1-563^:PMb256 interfaces as determined by the PISA server (https://www.ebi.ac.uk/pdbe/pisa/) and manual validation.

| Themis<sup>1-563</sup>:Grb2 |  |  | Themis<sup>1-563</sup>:PMb256 |  |  |
| --- | --- | --- | --- | --- | --- |
| Themis<sup>1-563</sup> | Grb2<sup>SH3C</sup> | Distance (Å) | Themis<sup>1-563</sup> | PMb256 | Distance (Å) |
| Hydrogen bonds |  |  | Hydrogen bonds |  |  |
| ARG 556 [NH1] | GLU 174 [OE1] | 3.07 | SER 462 [O] | TYR 32 [OH] | 3.88 |
| ARG 111 [NH1] | ARG 179 [O] | 3.60 | GLU 469 [O] | TYR 105 [N] | 2.81 |
| ARG 111 [NH2] | ARG 179 [O] | 3.74 | LEU 466 [O] | TYR 105 [OH] | 3.50 |
| ARG 111 [NH2] | ASP 181 [OD1] | 3.59 | GLU 469 [OE2] | TRP 108 [NE1] | 3.72 |
| ARG 180 [NH2] | ASP 181 [OD1] | 3.05 | ARG 464 [NH1] | ASP 99 [OD2] | 2.52 |
| TYR 541 [O] | GLN 162 [NE2] | 3.09 | ASP 471 [N] | VAL 103 [O] | 3.24 |
| GLU 547 [OE2] | ARG 179 [NH1] | 2.96 | ----- |  |  |
| GLU 483 [OE1] | ARG 179 [NH2] | 2.75 | Salt bridges |  |  |
| PRO 557 [O] | TRP 193 [NE1] | 3.56 | GLU 469 [OE1] | ARG 45 [NH2] | 3.94 |
| PRO 558 [O] | TRP 193 [NE2] | 3.58 | GLU 470 [OE1] | ARG 104 [NH1] | 3.99 |
| ASP 354 [OD2] | ARG 207 [NH2] | 2.45 | ARG 464 [NH1] | ASP 99 [OD2] | 2.52 |
| TYR 353 [OH] | ASN 208 [ND2] | 3.64 | ----- |  |  |
| PRO 554 [O] | TYR 209 [OH] | 3.13 | Buried residues (numerical order) |  |  |
| TYR 541 [OH] | ASN 214 [N] | 3.52 | VAL 463 | PHE 37 |  |
| ASP 204 [OD1] | ARG 215 [NH2] | 2.59 | ALA 474 | LYS 43 |  |
| PHE 205 [O] | ARG 215 [NH2] | 3.70 | ASN 497 | GLU 44 |  |
| SER 325 [O] | ASN 216 [ND2] | 3.55 | LEU 527 |  |  |
| ----- |  |  |  |  |  |
| Salt bridges |  |  |  |  |  |
| LYS 559 [NZ] | GLU 171 [OE1] | 3.44 |  |  |  |
| ARG 556 [NH1] | GLU 174 [OE1] | 3.07 |  |  |  |
| ARG 556 [NH1] | GLU 174 [OE2] | 3.65 |  |  |  |
| ARG 111 [NH2] | ASP 181 [OD1] | 3.59 |  |  |  |
| ARG 180 [NH2] | ASP 181 [OD1] | 3.05 |  |  |  |
| GLU 547 [OE2] | ARG 179 [NH1] | 2.96 |  |  |  |
| GLU 547 [OE2] | ARG 179 [NH2] | 3.68 |  |  |  |
| GLU 483 [OE1] | ARG 179 [NH2] | 2.75 |  |  |  |
| ASP 354 [OD2] | ARG 207 [NH1] | 3.53 |  |  |  |
| ASP 354 [OD1] | ARG 207 [NH2] | 3.63 |  |  |  |
| ASP 354 [OD2] | ARG 207 [NH2] | 2.45 |  |  |  |
| ASP 204 [OD2] | ARG 215 [NH2] | 3.48 |  |  |  |
| ASP 204 [OD1] | ARG 215 [NH2] | 2.59 |  |  |  |
| ----- |  |  |  |  |  |
| Buried residues (numerical order) |  |  |  |  |  |
| LEU 85 | TYR 160 |  |  |  |  |
| ILE 181 | PHE 165 |  |  |  |  |
| PHE 327 | PHE 182 |  |  |  |  |
| THR 351 |  |  |  |  |  |
| ARG 544 |  |  |  |  |  |
| ARG 545 |  |  |  |  |  |
| PRO 553 |  |  |  |  |  |

